# AdaFDR: a Fast, Powerful and Covariate-Adaptive Approach to Multiple Hypothesis Testing

**DOI:** 10.1101/496372

**Authors:** Martin J. Zhang, Fei Xia, James Zou

## Abstract

Multiple hypothesis testing is an essential component of modern data science. Its goal is to maximize the number of discoveries while controlling the fraction of false discoveries. In many settings, in addition to the p-value, additional information/covariates for each hypothesis are available. For example, in eQTL studies, each hypothesis tests the correlation between a variant and the expression of a gene. We also have additional covariates such as the location, conservation and chromatin status of the variant, which could inform how likely the association is to be due to noise. However, popular multiple hypothesis testing approaches, such as Benjamini-Hochberg procedure (BH) and independent hypothesis weighting (IHW), either ignore these covariates or assume the covariate to be univariate. We introduce `AdaFDR`, a fast and flexible method that adaptively learns the optimal p-value threshold from covariates to significantly improve detection power. On eQTL analysis of the GTEx data, `AdaFDR` discovers 32% and 27% more associations than BH and IHW, respectively, at the same false discovery rate. We prove that `AdaFDR` controls false discovery proportion, and show that it makes substantially more discoveries while controlling FDR in extensive experiments. `AdaFDR` is computationally efficient and can process more than 100 million hypotheses within an hour and allows multi-dimensional covariates with both numeric and categorical values. It also provides exploratory plots for the user to interpret how each covariate affects the significance of hypotheses, making it broadly useful across many applications.

## Introduction

Multiple hypothesis testing is an essential component in many modern data analysis workflows. A very common objective is to maximize the number of discoveries while controlling the fraction of false discoveries. For example, we may want to identify as many genes as possible that are differentially expressed between two populations such that less than, say, 10% of these identified genes are false positives.

In the standard setting, the data for each hypothesis is summarized by a p-value, with a smaller value presenting stronger evidence against the null hypothesis that there is no association. Commonly-used procedures such as Benjamini-Hochberg (BH)^1^ works solely with this list of p-values^2–6^. Despite being widely used, these multiple testing procedures fail to utilize additional information that is often available in modern applications that are not directly captured by the p-value.

For example, in expression quantitative trait loci (eQTL) mapping or genome-wide association studies (GWAS), single nucleotide polymorphism (SNP) in active chromatin state are more likely to be significantly associated with the phenotype^7^. Such chromatin information is readily available in public databases^8^, but is not used by standard multiple hypothesis testing procedures—it is sometimes used for post-hoc biological interpretation. Similarly, the location of the SNP, its conservation score, etc., can alter the likelihood for the SNP to be an eQTL. Together such additional information, called covariates, forms a feature representation of the hypothesis; this feature vector is ignored by the standard multiple hypothesis testing procedures.

In this paper, we present `AdaFDR`, a fast and flexible method that adaptively learns the decision threshold from covariates to significantly improve the detection power while having the false discovery proportion (FDP) controlled at a user-specified level. A schematic diagram for `AdaFDR` is shown in Figure 1. `AdaFDR` takes as input a list of hypotheses, each with a p-value and a covariate vector. Conventional methods like BH use only p-values and have the same p-value threshold for all hypotheses (Figure 1 top right). However, as illustrated in the bottom-left panel, the data may have an enrichment of small p-values for certain values of the covariate, which suggests an enrichment of alternative hypotheses around these covariate values. Intuitively, allocating more FDR budget to hypothesis with such covariates could increase the detection power. `AdaFDR` adaptively learns such pattern using both p-values and covariates, resulting in a covariate-dependent threshold that makes more discoveries under the same FDP constraint (Figure 1 bottom right).

**Figure 1.**
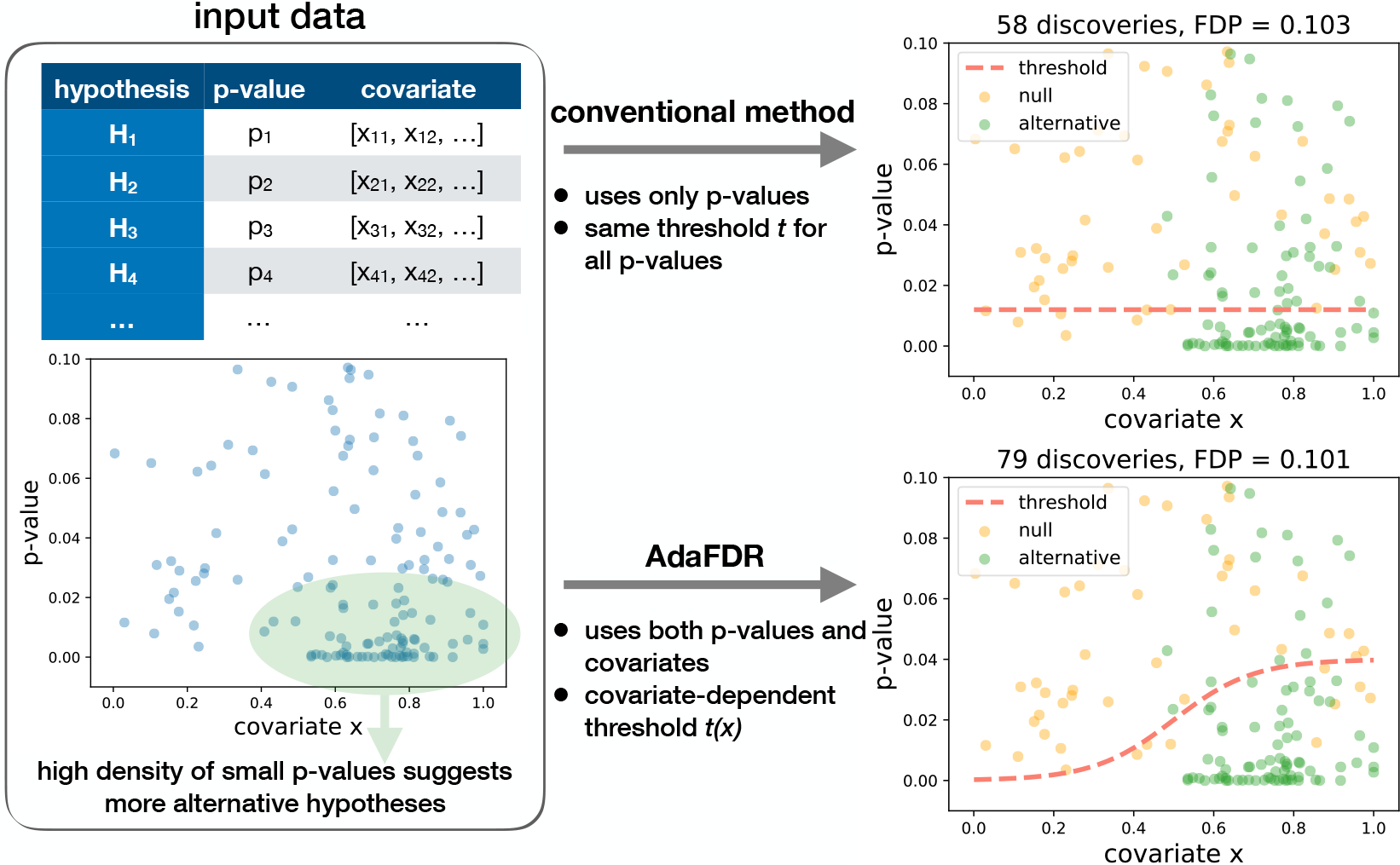
Intuition of `AdaFDR`. Top-left: As input, `AdaFDR` takes a list of hypotheses, each with a p-value and a covariate that could be multi-dimensional. Bottom-left: A toy example with a univariate covariate. The enrichment of small p-values in the bottom-right corner suggests that there are more alternative hypotheses there. Leveraging this structure can lead to more discoveries. Top-right: Conventional method uses only p-values and has the same p-value threshold for all hypotheses. Bottom-right: `AdaFDR` adaptively learns the uneven distribution of the alternative hypotheses, and makes more discoveries while controlling the false discovery proportion (FDP) at the desired level (0.1 in this case).

### Overview

`AdaFDR` extends conventional procedures like BH and Storey-BH (SBH)^2,3^ by considering multiple hypothesis testing with side information on the hypotheses. The input of `AdaFDR` is a set of hypotheses each with a p-value and a vector of covariates, whereas the output is a set of selected (also called rejected) hypotheses. For eQTL analysis, each hypothesis is one pair of SNP and gene, and the p-value tests for association between their values across samples. The covariate can be the location, conservation, and chromatin status at the SNP and the gene. The standard assumption of `AdaFDR` and all the related methods is that the covariates should not affect the p-values under the null hypothesis (see the Methods section for more discussion of this). `AdaFDR` learns the covariate-dependent p-value selection threshold by first fitting a mixture model using expectation maximization (EM) algorithm, where the mixtures model is a combination of a generalized linear model (GLM) and Gaussian mixture^9–11^. Then it makes local adjustments in the p-value threshold by optimizing for more discoveries. We prove that `AdaFDR` controls FDP under standard statistical assumptions in Theorem 1. `AdaFDR` is designed to be fast and flexible — it can simultaneously process more than 100 million hypotheses within an hour and allows multi-dimensional covariates with both numeric and categorical values. In addition, `AdaFDR` provides exploratory plots visualizing how each covariate is related to the significance of hypotheses, allowing users to interpret its findings. We also provide a much faster but slightly less powerful version, `AdaFDR-fast`, which uses only the EM step and skips the subsequent optimization. It can process more than 100 million hypotheses in around 5 minutes on a standard laptop.

We systematically evaluate the performance of `AdaFDR` across multiple datasets. We first consider the problem of eQTL discovery using the data from the Genotype-Tissue Expression (GTEx) project^7^. As covariates, we consider the distance between the SNP and the gene, the gene expression level, the alternative allele frequency as well as the chromatin states of the SNP. Across all 17 tissues considered in the study, `AdaFDR` has an improvement of 32% over BH and 27% over the state-of-art covariate-adaptive method independent hypothesis weighting (IHW)^12,13^. We next consider other applications, including three RNA-Seq datasets^14–16^ with the gene expression level as the covariate, two microbiome datasets^17,18^ with ubiquity (proportion of samples where the feature is detected) and the mean nonzero abundance as covariates, a proteomics datase^12,19^ with the peptides level as the covariate, and two fMRI datasets^20,21^ with the Brodmann area label^22^ as the covariate that represents different functional regions of human brain. In all experiments, `AdaFDR` shows a similar improvement. Finally, we perform extensive simulations, including ones from a very recent benchmark paper (Oct 31st 2018)^18^, to demonstrate that `AdaFDR` has the highest detection power while controlling the false discovery proportion (FDP) in various cases where the p-values may be either independent or dependent. The default parameters of `AdaFDR` are used for every experiment in this paper, both real data analysis and simulations, without any tuning. In addition to the experiments, we theoretically prove that `AdaFDR` controls FDP with high probability when the null p-values, conditional on the covariates, are independently distributed and stochastically greater than the uniform distribution, a standard assumption also made by related literature^13,23,24^.

### Related works

The problem of multiple hypothesis testing with covariates has recently been actively explored^12,13,24–28^. These works assume that for each hypothesis, we observe not only a p-value *P_i_* but also a general covariate **x**_*i*_ which is meant to capture the information on the significance of the hypothesis. However, the nature of this relationship is not known ahead of time and must be learned from the data. IHW^12,13^ groups the hypotheses into a pre-specified number of bins and applies a constant threshold for each bin to maximize the discoveries. It is practical, well-received by the community, and can scale up to 1 billion hypotheses. Yet it only supports the covariate to be univariate and uses a stepwise-constant function for the threshold, which limits its detection power. AdaPT^24^ cleverly uses a p-value masking procedure to control FDR. While IHW and `AdaFDR` need to split the hypotheses into multiple folds for FDR control, AdaPT can learn the threshold using virtually the entire data. However, such p-value masking procedure takes many iterations of optimization, and can be computationally expensive. Hence, while having high detection power, AdaPT usually takes a long time to run. `AdaFDR` is designed to achieve the best of both worlds: it has a speed comparable to IHW while using a flexible modeling strategy to have greater detection power than AdaPT.

There are also other methods in the field tailored for specific applications, where the domain knowledge can be used to increase the detection power. For example, gene set enrichment analysis (GSEA)^29^ uses the gene pathway information to identify classes of genes that are over-represented in a given set of genes. A recent work integrates genomic annotations into a Bayes hierarchical model to increase detection power in eQTL study^30^. Another incorporates phylogenetic tree information into a Bayesian model to increase the detection power in microbiome-wide multiple testing^31^. Compared to these methods, `AdaFDR` does not assume any prior knowledge about the covariates and learns the decision threshold in a completely data-driven manner. Hence, it is a more general approach that has a wider range of applications. Some other related works include non-adaptive p-value weighting^32–34,34,35^, estimation of the covariate-dependent null proportion^36–38^, and estimation of the local false discovery rate^39–43^.

`AdaFDR` is the mature development of and subsumes a previous, preliminary method that we called `NeuralFDR`^27^. Instead of using a neural network to model the discovery threshold as in `NeuralFDR`, `AdaFDR` uses a mixture model that lacks some flexibility but is much faster to optimize — for the GTEx data used in the `NeuralFDR` paper, it takes `NeuralFDR` 10+ hours to process but only 9 minutes for `AdaFDR`. Yet, `AdaFDR` maintains similar discovery power on the benchmark data used to test `NeuralFDR` (Supplementary Figure 3b). We systematically evaluated `AdaFDR` on many more settings and experiments than what was done for `NeuralFDR`.

## Results

### Discovering eQTLs in GTEx

We first consider detecting eQTLs using data from GTEx^7^. The GTEx project has collected both genetic variation data (SNPs) and gene expression data (RNA-Seq) from 44 human tissues, with sample sizes ranging from 70 (uterus) to 361 (muscle skeletal). Its goal is to study the associations between genotype and gene expression across humans. Each hypothesis test is to test if there is a significant association between a SNP and a gene, also referred to as an eQTL. A standard caveat is that a selected eQTL (either through small p-value or a FDR procedure) may not be a true *causal* SNP — it could tag a nearby causal SNP due to linkage disequilibrium. We should interpret the selected eQTLs with care; nonetheless, it is still valuable to discover candidate associations and local regions with strong associations while controlling FDR^12^.

We focus on cis-eQTLs where the SNP and the gene are close to each other on the genome (< 1 million base pairs). Previous works provide evidence that various covariates could be associated with the significance of cis-eQTLs^7,30,44,45^. In this study, we consider four covariates for each SNP-gene pair: 1) the distance from SNP to gene transcription start site (TSS); 2) the log10 gene expression level; 3) the alternative allele frequency (AAF) of the SNP; 4) the chromatin state of the SNP. Out of 44 tissues, we selected 17 whose chromatin state information is available^8^ and have more than 100 samples. For each tissue, p-values for all associations are tested simultaneously with numbers of hypotheses ranging from 140 to 180 million for different tissues, imposing a very-large-scale multiple hypothesis testing problem. We use a nominal FDR level of 0.01. Such experiments of testing all SNP-gene pairs simultaneously are also performed in^12,13^. An alternative analysis workflow is to first discover significant genes (eGenes) and then match significant SNPs (eVariants) for each eGene^30^.

As shown in Figure 2a, `AdaFDR` and its fast version consistently make more discoveries than other methods in every tissue. On average, it has an improvement of 32% over BH and 27% over IHW. Next we investigate whether using the eQTL p-values of an existing tissue could boost the power of discovering eQTLs in a new tissue. To simulate this scenario, we consider specifically the two adipose tissues, Adipose_Subcutaneous and Adipose_Visceral_Omentum. For each of them, we use the −log10 p-values from the other tissue as an additional covariate— e.g. for Adipose_Subcutaneous, the −log10 p-value of Adipose_Visceral_Omentum is used as an extra covariate. Leveraging previous eQTL results substantially increases discovery power (Figure 2b); the p-value augmentation (AdaFDR (aug)) yields 56% and 83% more discoveries for the two adipose tissues compared to BH. We then perform a control experiment, where the augmented p-values, instead of coming from the other adipose tissue that is similar to the one under investigation, are from a brain tissue (Brain_Caudate_basal_ganglia) that is very different from the adipose tissue (Figure 2 in the GTEx paper^7^). In this case, the improvement in the number of discoveries due to the extra covariate vanishes for the two tissues (AdaFDR (ctrl), which is consistent with the idea that `AdaFDR` learns to leverage shared genetic architecture in closely related tissues to improve power. This analysis suggests that we can potentially greatly improve eQTL discovery by leveraging related tissues during multiple hypothesis testing. We provide additional supporting experiments for the two colon tissues in Supplementary Figure 1a.

**Figure 2.**
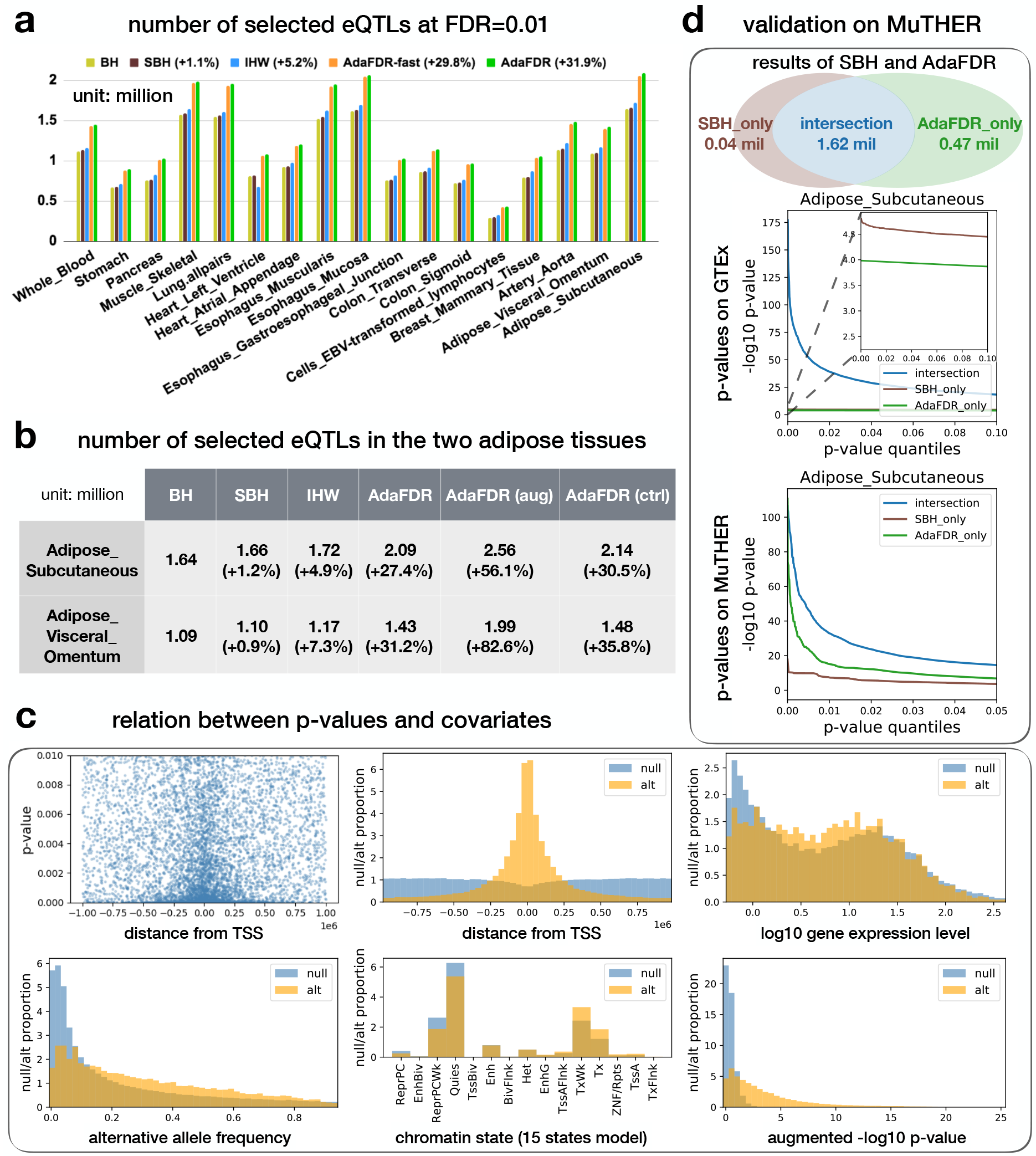
Analysis of the GTEx data. (a) Results of 17 tissues considered in the study. `AdaFDR` and its fast version consistently make more discoveries than other methods. (b) Results on the two adipose tissues where the −log10 p-value from another tissue was added as an extra covariate. Using p-values from a similar tissue (AdaFDR (aug)) yields significantly more discoveries than using p-values from an unrelated tissue (AdaFDR (ctrl)). (c) Top-left: P-values (y-axis) plotted against the distances from TSS (x-axis); each dot corresponds to one SNP-gene pair. Small p-values at the center suggest that hypotheses with smaller distances from TSS are more likely to be significant. Other panels: `AdaFDR`-estimated null hypothesis distribution (blue) and alternative hypothesis distribution (orange) with respect to each covariate. Higher values of the orange distribution suggest an enrichment of alternative hypotheses. (d) Top: Discoveries made by SBH and `AdaFDR`. Middle: The p-values of these discoveries—SBH-only p-values are smaller than `AdaFDR`-only p-values on GTEx. Bottom: The p-values of the same set of discoveries on the independent MuTHER data, where `AdaFDR`-only p-values are smaller than SBH-only p-values, suggesting that `AdaFDR`-only discoveries are more likely to be true discoveries.

`AdaFDR` also characterizes how each covariate affect the significance level of the hypotheses. The results for Adi-pose_Subcutaneous are shown in Figure 2c as an example. We first consider the distance from TSS and the top-left panel provides a simple visualization, where for each hypothesis (downsampled to 10k), the p-values are plotted against the distances from TSS. There is a strong enrichment of small p-values when the distance is close to 0, indicating that the SNP and gene are more likely to have a significant association if they are close to each other. In the top-center panel, `AdaFDR` characterizes such relationship by providing estimates of the null hypothesis distribution (blue) and the alternative hypothesis distribution (orange), with respect to the distance from TSS. It learns that an alternative hypothesis is more likely to appear at the center, where the distance from TSS is small, consistent with previous works^7,44^.

`AdaFDR` interprets other covariates in a similar fashion. Figure 2c top-right panel indicates that genes with higher expression levels are more likely to have significant associations, in agreement with previous observations^12,24^. SNPs with AAF close to 0.5 are also more likely to have significant associations. In addition, the bottom-center panel indicates that SNPs with active chromatin states—Tx (strong transcription), TxWk (weak transcription), TssA (active TSS)—are more likely to have significant associations as compared to SNPs with inactive states—Quies (quiescent), ReprPC (repressed PolyComb) ReprPCWk (weak repressed PolyComb). Finally, the bottom-right panel shows that p-values from the augmented tissue Adipose_Visceral_Omentum are positively correlated with the significance of the associations. See Supplementary Figure 1b for analogous results on the Colon_Sigmoid tissue.

We use adipose eQTL data from the Multiple Tissue Human Expression Resource (MuTHER) project^46^ to validate our GTEx eQTL discoveries. The participants in MuTHER are disjoint from the GTEx participants, making MuTHER an independent datasets. For this analysis, we compare the testing results of `AdaFDR` on Adipose_Subcutaneous with that of Storey-BH (SBH), which is known to be a better baseline than BH. As shown in the top panel of Figure 2d, `AdaFDR` detects almost all discoveries made by SBH while having 26% more discoveries. The p-values of these discoveries are shown in the middle panel of Figure 2d, where x-axis is the p-value quantile and y-axis is the −log10 p-value. Hypotheses discovered by both methods have significantly smaller GTEx p-values while SBH-only p-values are smaller than AdaFDR-only p-values in the GTEx data; the latter is due to the fact that SBH uses the same threshold for all p-values. On the MuTHER validation data, the eQTLs discovered only by `AdaFDR` have more significant p-values than the eQTLs discovered only by SBH. This reveals a counter-intuitive behaviour of `AdaFDR`: it rejects some hypotheses with larger p-values if these SNPs have covariates that indicate a higher likelihood of eQTL. The MuTHER data validates this strategy—`AdaFDR` is able to discover more eQTLs on GTEx and the discovered eQTLs have more significant replication results on MuTHER.

`AdaFDR` can be broadly applied to any multiple testing problem where we have covariates for the hypotheses. This includes many highthroughput biological studies beyond eQTL. Here we evaluate its applications to RNA-seq, microbiome, proteomics and fMRI imaging data. In all cases, `AdaFDR` significantly outperforms current state-of-the-art methods.

### Small GTEx data

AdaPT cannot be run on the full GTEx data due to its computational limitations. In order to perform a direct comparison between `AdaFDR` and AdaPT, we created a small GTEx data that contains the first 300k associations from chromosome 21 for the two adipose tissues. Even this small data takes AdaPT around 15 hours to process compared to less than 20 minutes for other methods. As shown in Figure 3a, `AdaFDR` has most number of discoveries in both experiments while AdaPT has slightly less. In addition, all covariate-adaptive methods have significant improvement over the non-adaptive methods (BH, SBH).

**Figure 3.**
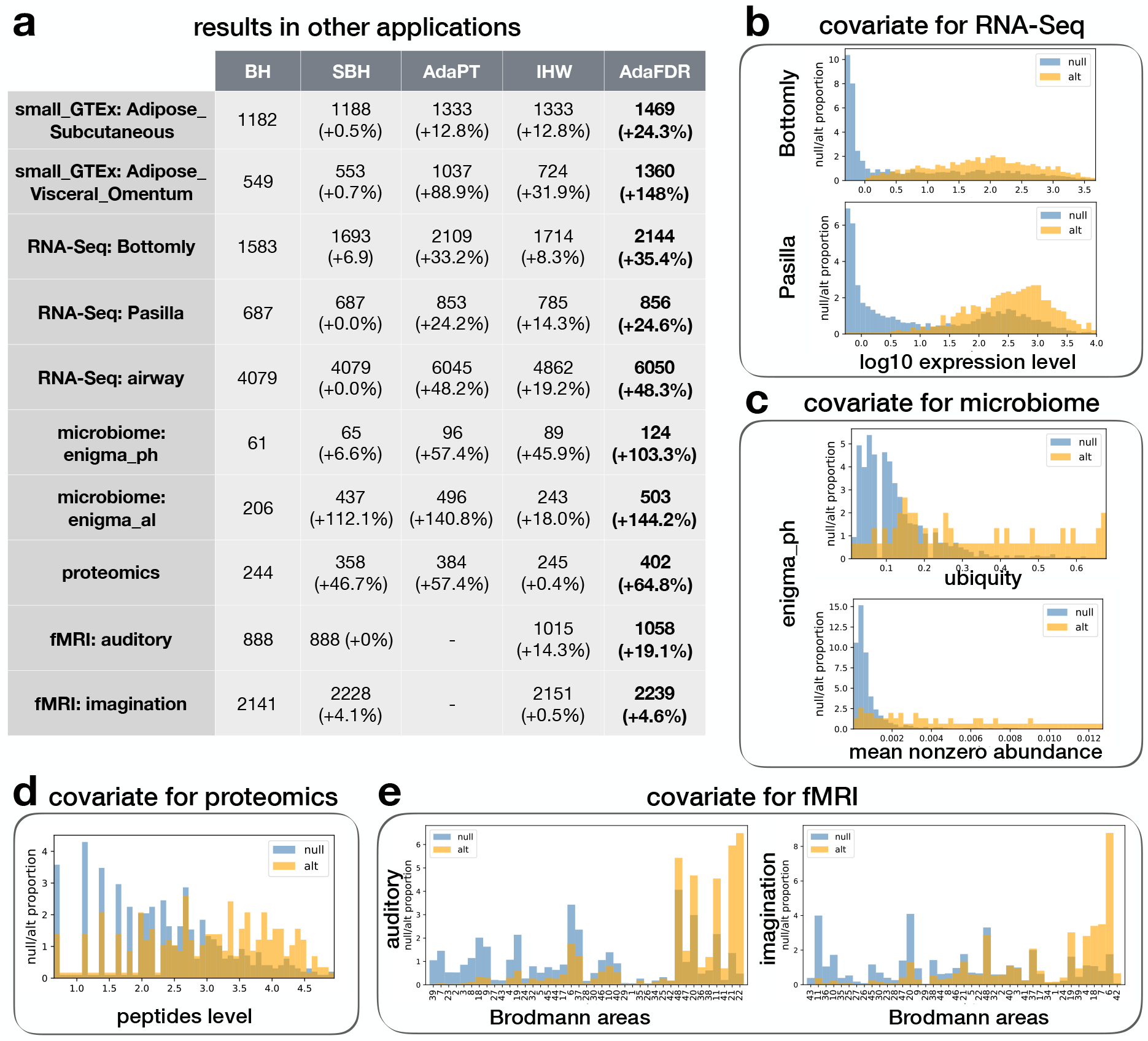
(a) The number of discoveries of various methods on two small GTEx eQTL datasets, three RNA-Seq differential expression datasets, two microbiome datasets, one proteomics dataset, and two fMRI datasets. The fMRI results for AdaPT are omitted since the AdaPT software does not support categorical covariates. (b) Covariate visualization for RNA-Seq datasets. (c) Covariate visualization for microbiome dataset. (d) Covariate visualization for proteomics dataset. (e) Covariate visualization for fMRI datasets.

### RNA-Seq data

We considered three RNA-Seq datasets that were used for differential expression analysis in AdaPT and IHW, i.e. the Bottomly data^15^, the Pasilla data^16^ and the airway data^14^. Here, the log expression level is used as the covariate, and the FDR level is set to be 0.1. The results are shown in Figure 3a, where `AdaFDR` and AdaPT have a similar number of discoveries (`AdaFDR` is consistently higher), and both are substantially more powerful than others. All covariate-adaptive methods make significantly more discoveries than the non-adaptive methods. In addition, the covariate patterns learned by `AdaFDR` are shown in Figure 3b for the Bottomly data and the Pasilla data, and in Supplementary Figure 2c for the airway data. The alternative hypotheses are more likely to occur when the expression levels are high, consistent with previous findings^12,13,24^.

### Microbiome data

We considered a subset of microbiome data from the Ecosystems and Networks Integrated with Genes and Molecular Assemblies (ENIGMA), where samples were acquired from monitoring wells in a site contaminated by former waste disposal ponds and all sampled wells have various geochemical and physical measurements^17,18^. Following the original study, we performed two experiments to test for correlations between the operational taxonomic units (OTUs) and the pH, Al respectively. Ubiquity and the mean nonzero abundance are used are covariates, where the ubiquity is defined as the proportion of samples in which the OTU is present. The FDR level is set to be 0.2 for more discoveries and the fast version of `AdaFDR` is used due to the small sample size. As shown in Figure 3a, `AdaFDR` is significantly more powerful than other methods. The covariates are visualized in Figure 3c for the pH test and Supplementary Figure 2b for the Al test. The alternative hypotheses are more likely to occur when both the ubiquity and the mean nonzero abundance are high. This may be because that a higher level of these two quantities improves the detection power similar to the expression level in the RNA-Seq case.

### Proteomics data

We considered a proteomics datasets where yeast cells treated with rapamycin were compared to yeast cells treated with dimethyl sulfoxide (2 × 6 biological replicates)^12,19^. Differential abundance of 2,666 proteins is evaluated using Welch’s t-test. The total number of peptides is used as covariate that is quantified across all samples for each protein. The FDR level is set to be 0. 1 and the fast version of `AdaFDR` is used due to the small sample size. As shown in Figure 3a, `AdaFDR` is significantly more powerful than other methods. The covariate is visualized in Figure 3d where a higher level of peptides increases the likelihood for the alternative hypotheses to occur. This is expected since the peptides level is similar to the expression level in the RNA-Seq data.

### fMRI data

We considered two functional magnetic resonance imaging (fMRI) experiments where the human brain is divided spatially into isotropic voxels and the null hypothesis for each voxel is that there is no response to the stimulus^20^. The first experiment was done on a single participant with auditory stimulus and the second was done on a healthy adult female participant where the stimulus was to ask the person to imagine playing tennis^21^. We use the Brodmann area label, which represents different functional regions of the human brain^22^, as covariate for each voxel. The FDR level is set to be 0.1 and the fast version of `AdaFDR` is used due to the inflation of p-values at 1. As shown in Figure 3a, `AdaFDR` is significantly more powerful than other methods. The result of AdaPT is omitted since it does not support categorical covariates, and directly running the GAM model yields a result much worse than BH. The covariate is visualized in Figure 3e. For the auditory experiment, the Brodmann areas corresponding to auditory cortices, namely 41, 42, 22, are among areas where the alternative hypotheses are most likely to occur. For the tennis imagination experiment, multiple cortices seem to respond to this stimulus, including auditory cortex (42), visual cortices (18,19), and motor cortices (4,6,7).

### Simulation studies

In order to systematically quantify the FDP and power of all the methods, we conducted extensive analysis of synthetic data where we know the ground truth. Each experiment is repeated 10 times and 95% confidence intervals are provided. In Figure 4a, the top two panels correspond to a simulated data with one covariate while the bottom two panels correspond to a simulated data with weakly-dependent p-values generated according to a previous paper^4^. In both simulations, all methods control FDR while `AdaFDR` has significantly larger power. Additional simulation experiments with strongly-dependent p-values and higher dimensional covariates can be found in Supplementary Figure 4a, where similar results are observed. Detailed descriptions of the synthetic data can be found in Supplementary Section 3.

**Figure 4.**
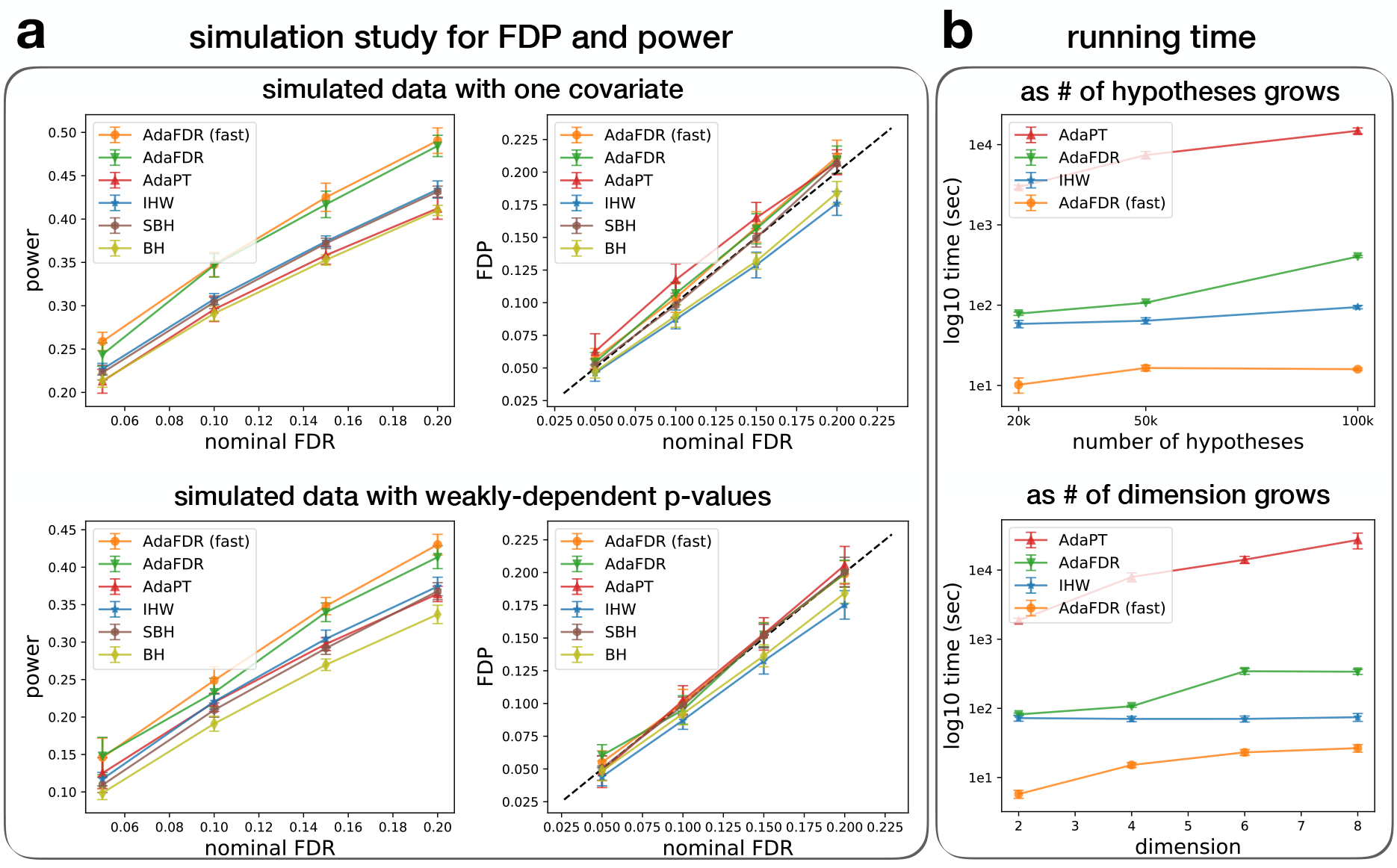
(a) The number of discoveries of various methods on three RNA-Seq differential expression datasets and two small GTEx eQTL datasets. (b) Covariate visualization for RNA-Seq datasets. Note that the AdaPT software does not support categorical covariates and can not be ran on the two fMRI datasets. (c) Simulation of FDP and power on an independent case (top) and a weakly-dependent case (bottom). (d) Running time analysis. Top: as the number of hypotheses grows. Bottom: as the covariate dimension grows.

We also investigate the running time of different methods. In Figure 4b, all experiments are repeated 5 times and the 95% confidence intervals are provided. The top panel uses a simulated datasets with 2d covariate, with the number of hypotheses varying from 20k to 100k. `AdaFDR-fast` takes 10s to run while both `AdaFDR` and IHW finished within a reasonable time of around 100s. AdaPT, however, needs a few hours to finish, significantly slower than other methods. In the bottom panel, the number of hypotheses is fixed to be 50k and the covariate dimension varies from 2 to 8; a similar result is observed.

After we have finished our initial paper, a very recent work^18^ on the bioRxiv (October 31, 2018) proposed a new set of benchmark experiments to compare state-of-the-art multiple testing methods including IHW, AdaPT and an additional method of Boca-Leek (BL) that is on the bioRxiv^47^. We use their main simulation benchmark that includes two RNA-Seq *in silico* experiments, one experiment with uninformative covariate, and another two experiments that vary the number of hypotheses and the null proportion respectively. We run `AdaFDR` on this benchmark without any modification or tuning; `AdaFDR` achieves greater power than all other methods while controlling FDR (Supplementary Figure 4, 5). `AdaFDR` reduces to SBH when the covariate is not informative, indicating that it is not overfitting the uninformative covariate (Supplementary Figure 4e).

## Discussion

Here we propose AdaFDR, a fast and flexible method that efficiently utilizes covariate information to increase detection power. Extensive experiments show that `AdaFDR` has greater power than existing approaches while controlling FDR. We discuss some of its characteristics and limitations.

Our theory proves that `AdaFDR` controls FDP in the setting when the null hypotheses are independent (the alternative hypotheses can have arbitrary correlations, see Theorem 1). This is a standard assumption also used in BH, SBH, IHW and AdaPT. To check the robustness of `AdaFDR` when there is model mismatch, we have performed systematic simulations with different p-value correlation structures to demonstrate that `AdaFDR` still controls FDP even when the null hypotheses are not independent. Moreover, although there are correlations among SNPs in the eQTL study, we show that the discoveries made by `AdaFDR` on the GTEX data replicate well on the independent MuTHER data with a different cohort. These suggest that `AdaFDR` behaves well when there is a dependency between null p-values. Since none of the other methods popular methods—BH, SBH, IHW, AdaPT—provides FDR control under arbitrary dependency, our comparison experiments are fair. `AdaFDR` can potentially be extended to allow arbitrary dependency using a similar idea as discussed in IHW^13^. Specifically, hypotheses should be split in such a way that the p-values from the two folds are independent, though they may have dependency within each fold. As a result, the learned threshold is independent of the fold it is applied onto. Then ideas discussed in the Benjamini-Yekutieli paper^6^ can be used to scale the threshold to allow arbitrary dependency^13^.

The typical use-case for `AdaFDR` is when there are many hypotheses to be tested simultaneously — ideally more than 10k. This is because `AdaFDR` needs many data to learn the covariate-adaptive threshold and to have an accurate estimate of FDP. A similar recommendation on the number of hypotheses is also made for IHW. When we have a smaller number of hypotheses, the discoveries are still valid but need to be treated with precaution — ideally with some orthogonal validations.

The scalability of `AdaFDR` and its ability to handle multivariate discrete and continuous covariates makes it broadly applicable to any multiple testing applications where additional information is available. While we focus on genomics experiments in this paper—because most of the previous methods were also evaluated on genomics experiments — it would be interesting to also apply `AdaFDR` to other domains such as imaging association analysis.

## Methods

### Definitions and notations

Suppose we have *N* hypothesis tests and each of them can be characterized by a p-value *P_i_,* a *d*-dimensional covariate **x**_*i*_, and a indicator variable *h_i_* with *h_i_* = 1 representing the hypothesis to be true alternative. Then the set of true null hypotheses *ℋ*_0_ and the set of true alternative hypotheses ℋ_1_ can be written as ℋ_0_ = {*i*: *i* ∈ [*N*], *h_i_* = 0} and ℋ_1_ = {*i*: *i* ∈ [*N*], *h_i_* = 1}, where we adopt the notation 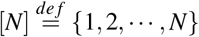. Given a threshold function *t*(**x**), we reject the *it*th null hypothesis if *P_i_* ≤ *t*(**x_i_**). The number of discoveries D(*t*) and the number of false discoveries FD(*t*) can be written as 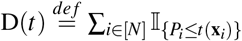 and 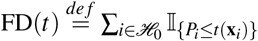. The false discovery proportion (FDP) is defined as 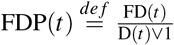, where 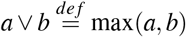. The expected value of FDP is the false discovery rate 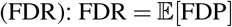^23^.

### Multiple testing via AdaFDR

`AdaFDR` can take as input multi-dimensional covariates **x**. The key assumption is that the null p-values remain uniform regardless of the covariate value while others, including the alternative p-values and the likelihood for the hypotheses to be true null/alternative, may have arbitrary dependencies on the covariate. This is a standard assumption in the literatur^12,23,24^. For example, in the case of AAF, the null p-values are uniformly distributed independent of AAF since the gene expression has no association with the SNP under the null hypothesis. However, the alternative p-values may depend on AAF since the associations are easier to detect/yield smaller p-values if the AAF is close to 0.5.

`AdaFDR` aims to optimize over a set of decision rules 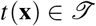 to maximize the number of discoveries, subject to the constraint that the FDP is less than a user-specified nominal level *α*. Conceptually, this optimization problem can be written as

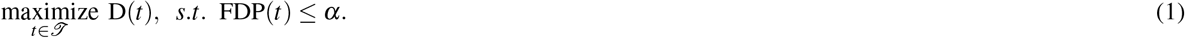

There are three challenges in this optimization problem: 1. the set of decision thresholds 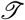 needs to be parameterized in such a way that both captures the covariate information and scales well with the covariate dimension; 2. the actual FDP is not directly available from the data; 3. direct optimization of (1) may cause overfitting and hence lose FDR control.

For the first challenge, intuitively, the decision threshold should have large values where the alternative hypotheses are enriched. Such enrichment pattern, as discussed the `NeuralFDR` paper^27^, usually consists of local “bumps” at certain covariate locations and a global “slope” that represents generic monotonic relationships. For example, the distance from TSS and the AAF in Figure 2c correspond to the bump structure (at 0 and 0.5 respectively) whereas the rest of the covariates correspond to the slope structure. `AdaFDR` addresses these two structures by using a mixture of generalized linear model (GLM) and *K*-component Gaussian mixture (with diagonal covariance matrices), i.e.,

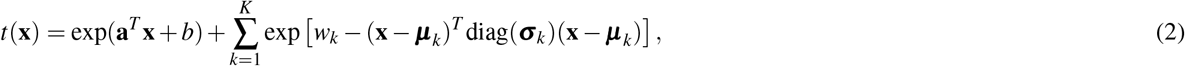

where diag(**σ**_*k*_) represents a diagonal matrix with diagonal elements specified by the d-dimensional vector **σ**_*k*_. The set of parameters to optimize can be written as 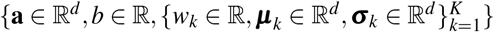. We choose to use the diagonal covariance matrices for Gaussian mixture to speed up the optimization. As a result, the number of parameters grows linearly with respect to the covariate dimension *d*, and the parameters can be easily initialized via EM algorithm, as described below.

For the second challenge, we use a “mirror estimator” to estimate the number of false discoveries of a given threshold function *t*,

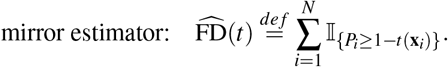

Such estimator has been used in recent work^24,27,48,49^ and yields a conservative estimate of the true number of false discoveries (FD), in the sense that its expected value is larger than that of the true FD under mild assumptions (Lemma 1 in Supplementary Materials). Furthermore, FDP can be simply estimated as 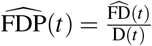.

For the third challenge, `AdaFDR` controls FDP with high probability via hypothesis splitting. The hypotheses are randomly split into two folds; a separate decision threshold is learned on each fold and applied on the other. Since the learned threshold does not depend on the fold of data onto which it is applied, FDP can be controlled with high probability — such statement is made formal in Theorem 1. We note that in multiple testing by `AdaFDR`, the learning-and-testing process is repeated twice, with each fold being the training set at one time and the testing set at the other. Figure 5 shows one of such process with fold 1 being the training set.

**Figure 5.**
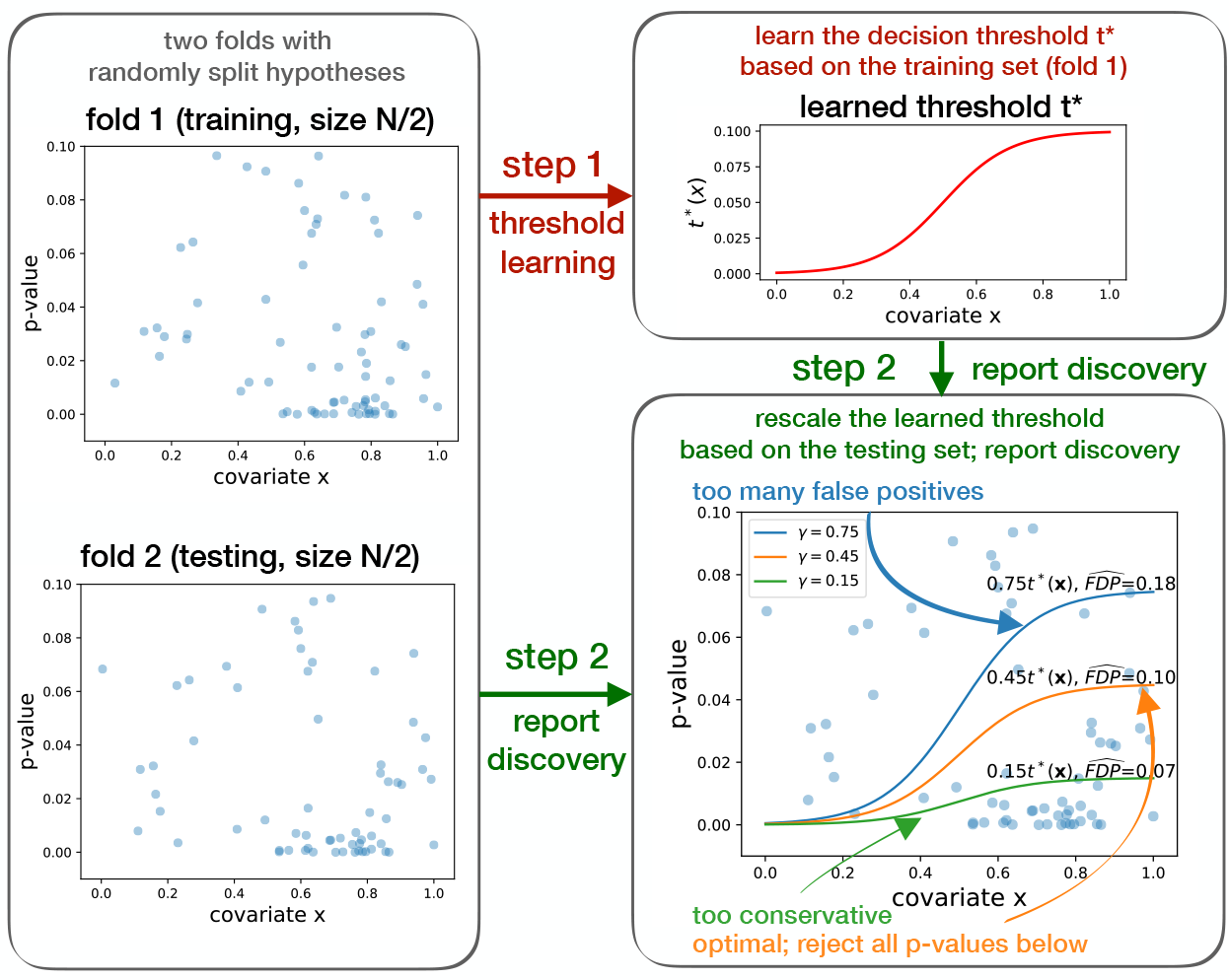
Schematic of the `AdaFDR` learning and testing process. Fold 1 is the training set and fold 2 is the testing set (left panel). In step 1, a decision threshold *t*\*(x) is learned on the training set via solving the optimization problem (1) (upper-right panel). In step 2, as shown in the bottom-right panel, this learned threshold *t*\*(x) is first rescaled by a factor γ*, defined as the largest number whose corresponding mirror-estimated FDP on the testing set is less than *α* (orange). Then all p-values on the testing set below the rescaled threshold are rejected. Here the nominal FDP is *α* = 0.1.

The full algorithm is described in Algorithm 1. Here, for example, D_*train*_(*t*), D_*test*_(*t*) are understood as the number of discoveries on the training set and the testing set respectively. Similar notations are used for other quantities like FDP(*t*) and the mirror estimate 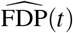 without explicit definition.

`AdaFDR` follows a similar strategy as our preliminary work `NeuralFDR`^27^, which it subsumes: both methods use the mirror estimator to estimate FDP and use hypothesis splitting for FDP control. The main difference is on the modeling of the decision threshold *t*: `NeuralFDR` uses a neural network, which is flexible enough but hard to optimize. `AdaFDR`, in contrast, adopts the simpler mixture model that may lack certain flexibility but is much easier to optimize. This change of modeling, however, does not seems to reduce much of the detection power for `AdaFDR`. As shown in Supplementary Figure 3b, the performance of `AdaFDR` is similar to that of `NeuralFDR`, while `AdaFDR` is orders of magnitude faster.

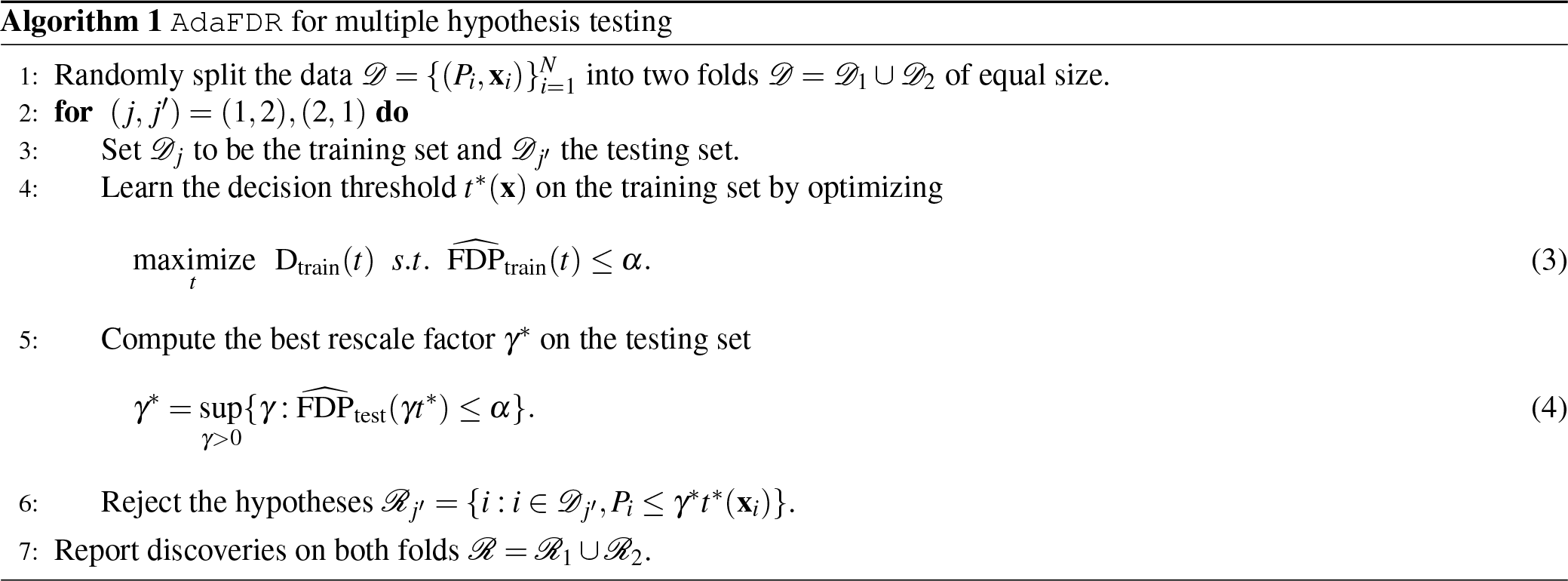

### Optimization

Recall that the optimization is done solely on the training set 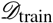. Substituting FDP in (1) with its mirror estimate we can rewrite the optimization problem as

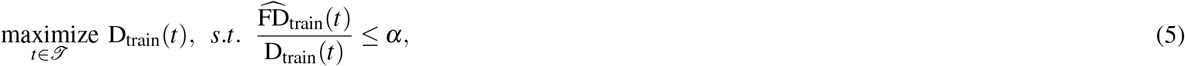

where 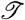, the set of decision thresholds to optimize over, corresponds to the mixture model (2). Our strategy is to first compute a good initialization point and then perform optimization by gradient descent on a relaxed problem. We note that a better solution to the optimization problem will give a better detection power. However, the FDP control guarantee holds *regardless* of the decision threshold we come up with.

- **Initialization**: Let *π*_0_(**x**) and *π*_1_ (**x**) be the distributions for the null hypotheses and the alternative hypotheses, over the covariate **x**, respectively. Following the intuition that the threshold *t* (**x**) should be large when the number of alternative hypotheses is high and the number of null hypotheses is low, it is a good heuristic to let

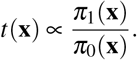 This is done in `AdaFDR` as follows. First, covariates with p-values larger than 0.75, i.e. 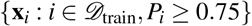, are treated as an approximate ensemble of the null hypotheses, and those with p-values smaller than the BH threshold, i.e. 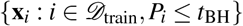, are treated as an approximate ensemble of the alternative hypotheses. Then first, a mixture model same as (2) is fitted on the null ensemble 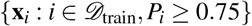 using EM algorithm, resulting in an estimate of the null hypothesis distribution 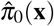. Second, each point in the alternative ensemble 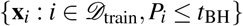 receives a sample weight 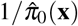. Last, the mixture model (2) is fitted on the weighted alternative ensemble using EM algorithm to obtain the final initialization threshold. The details of the EM algorithm can be found in Supplementary SubSection 2.3.
- **Optimization**: First, a Lagrangian multiplier is used to deal with the constraint:

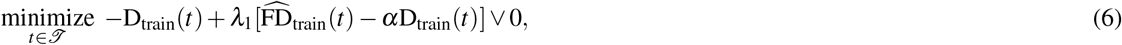

where λ_1_ is chosen heuristically to be 10/α. Second, the sigmoid function is used to deal with the discontinuity of the indicator functions in D_train_(*t*) and 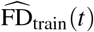:

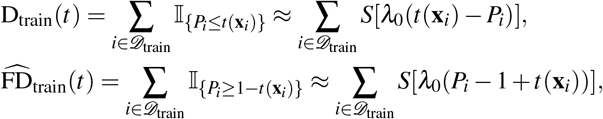

where 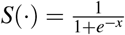 is the sigmoid function and λ_0_ is automatically chosen at the beginning of the optimization such that the smoothed versions are good approximations to the original ones. Finally, the Adam optimizer^50^ is used for gradient descent.

### FDP control

We would like to point out that the mirror estimate is more accurate when its value is large. Hence, when the number of rejections is small (<100), the result should be treated with precaution. However, this should not be a major concern since in the target applications of `AdaFDR`, usually thousands to millions of hypotheses are tested simultaneously, and hundreds to thousands of hypotheses are rejected. In those cases, the mirror estimate is accurate. Hence, we further require that for each fold, the best scale factor *γ*\* should have a number of discoveries exceeding *c*_0_*N* for some pre-specified small proportion *c*_0_; failing to satisfy this condition will result in no rejection in this fold. In other words, we consider a modified version of Alg. 1 with (4) substituted by setting

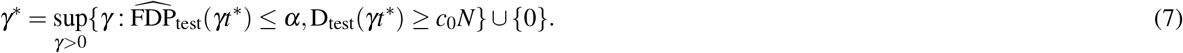

Our FDP control on this modified version can be stated as follows.

#### Theorem 1.

*(FDP control) Assume that all null p-values P_i_* ∈ ℋ_0_, *conditional on the covariates, are independently and identically distributed (i.i.d.) following* Unif[0,1]. *Then with probability at least* 1-*δ*, *`AdaFDR` with the modification* (7) *controls FDP at level* (1 + *ε*)*α*, *where* 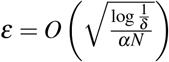.

The assumption made in Theorem 1 is standard in the literature^13,24^ and can be easily relaxed to the assumption that the null p-values, conditional on the covariates, are independently distributed and stochastically greater than Unif[0,1] (Supplementary SubSection 4.1). In addition, Theorem 1 is strictly stronger than the one for `NeuralFDR` (Supplementary SubSection 2.2).

### Covariate visualization via AdaFDR_explored

`AdaFDR` also provides a `FeatureExplore` function that can visualize the relationship between each covariate and the significance of hypotheses, in terms of estimated distributions for the null hypothesis and the alternative hypothesis with respect to each covariate, as those shown in Figure 2c and Figure 4b. This is done as follows. First, for the entire datasets, covariates with p-values greater than 0.75, i.e. {**x**_*i*_: *i* ∈ [*N*], *P_i_* ≥ 0.75}, are treated as an approximate ensemble of the null hypotheses, and those with p-values less than the BH threshold, i.e. {**x**_*i*_: *i* ∈ [*N*], *P_i_* ≤ *t*_BH_}, are treated as an approximate ensemble of the alternative hypotheses. Then, the null hypothesis distribution and the alternative hypothesis distribution are estimated from these two ensembles using kernel density estimation (KDE) for continuous covariates and simple count estimator for categorical covariates. In addition, for categorical covariates, the categories are reordered based on the ratio between the estimated alternative probability and null probability 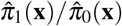.

## Acknowledgements

We would like to thank David Tse, Liuhua Lei, Nikolaos Ignatiadis and Vivek Bagaria for helpful discussions. MZ and FX are partially supported by Stanford Graduate Fellowship. JZ is supported by the Chan-Zuckerberg Initiative and National Science Foundation (NSF) Grant CRII 1657155.

## Author contributions statement

MZ designed the algorithm and conducted the experiments with the help of FX. MZ performed the theoretical analysis. MZ and JZ wrote the manuscript. JZ supervised the research. All authors reviewed the manuscript.

## Additional information

### Code availability

- The code for the paper is available at https://github.com/martinjzhang/AdaFDRpaper
- The software is available at https://github.com/martinjzhang/adafdr

#### Data availability

The GTEx data for the two adipose tissues and all other data are deposited into the online repository. See the github repository `AdaFDRpaper` for more information.

## Supplemental Materials

### 1 Additional Results

**Supplementary Figure 1.**
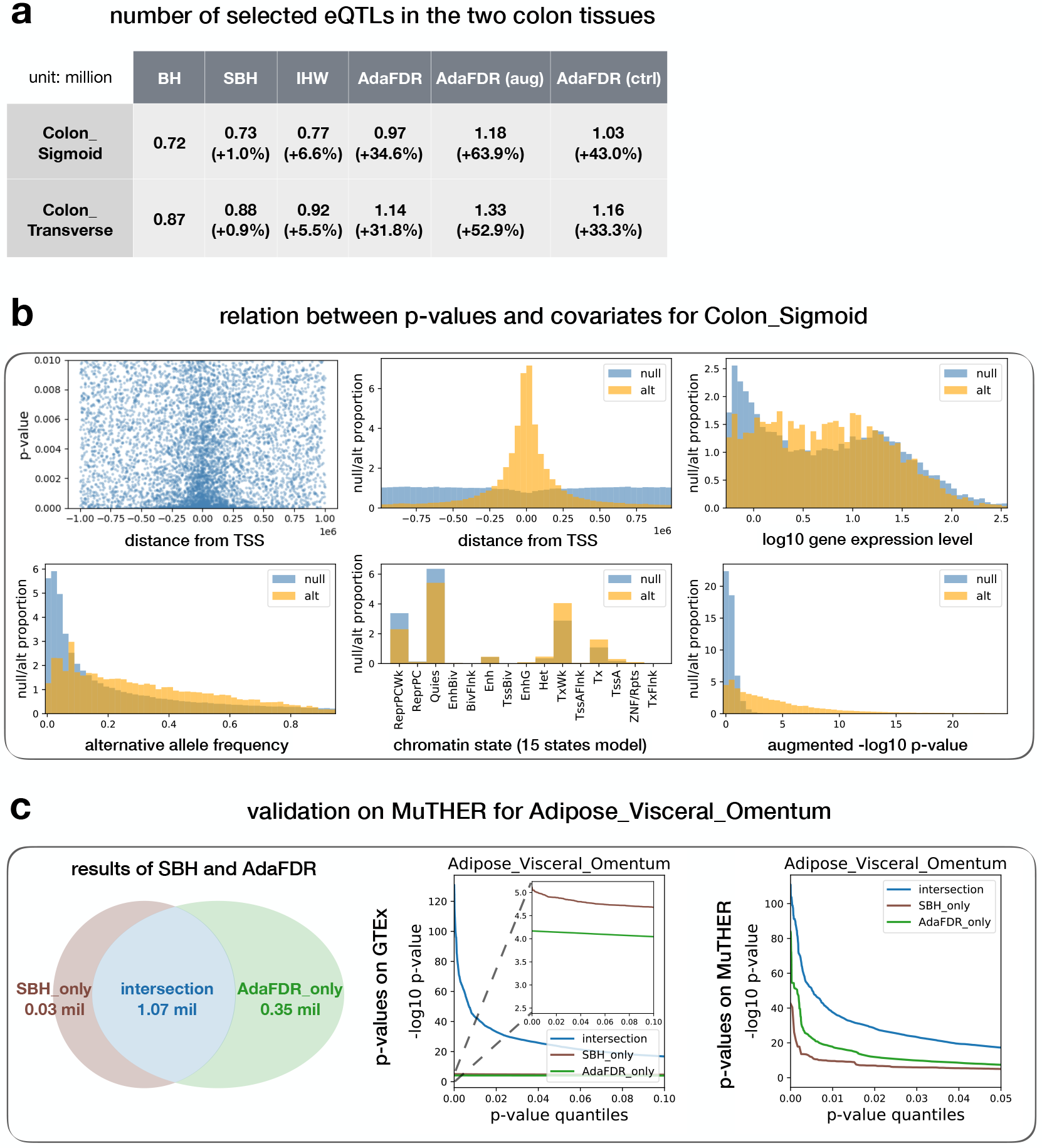
Additional results on the GTEx data. (a) Results on the two colon tissues. (b) Feature visualization for Colon_Sigmoid (c) Validation on MuTHER for Adipose_Visceral_Omentum.

**Supplementary Figure 2.**
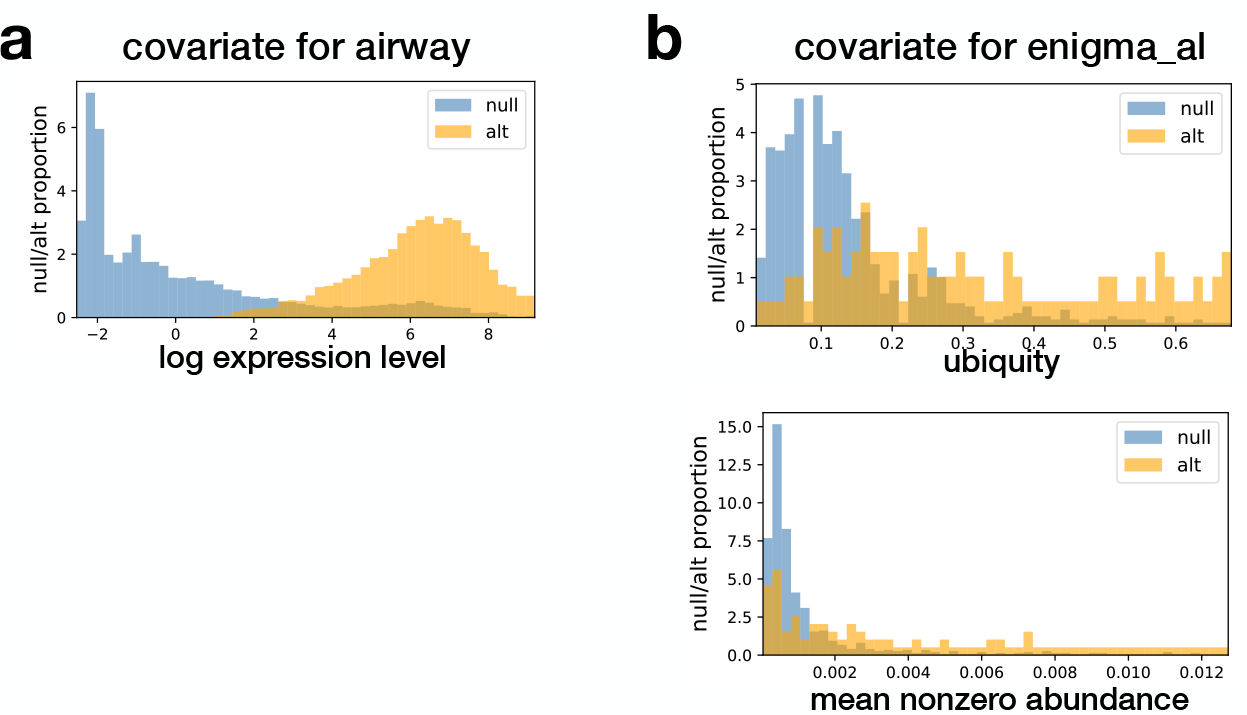
(a) The covariate visualization for the RNA-Seq airway data. (b) The covariate visualization for the microbiome enigma_al data. Top: ubiquity; bottom: mean nonzero abundance.

**Supplementary Figure 3.**
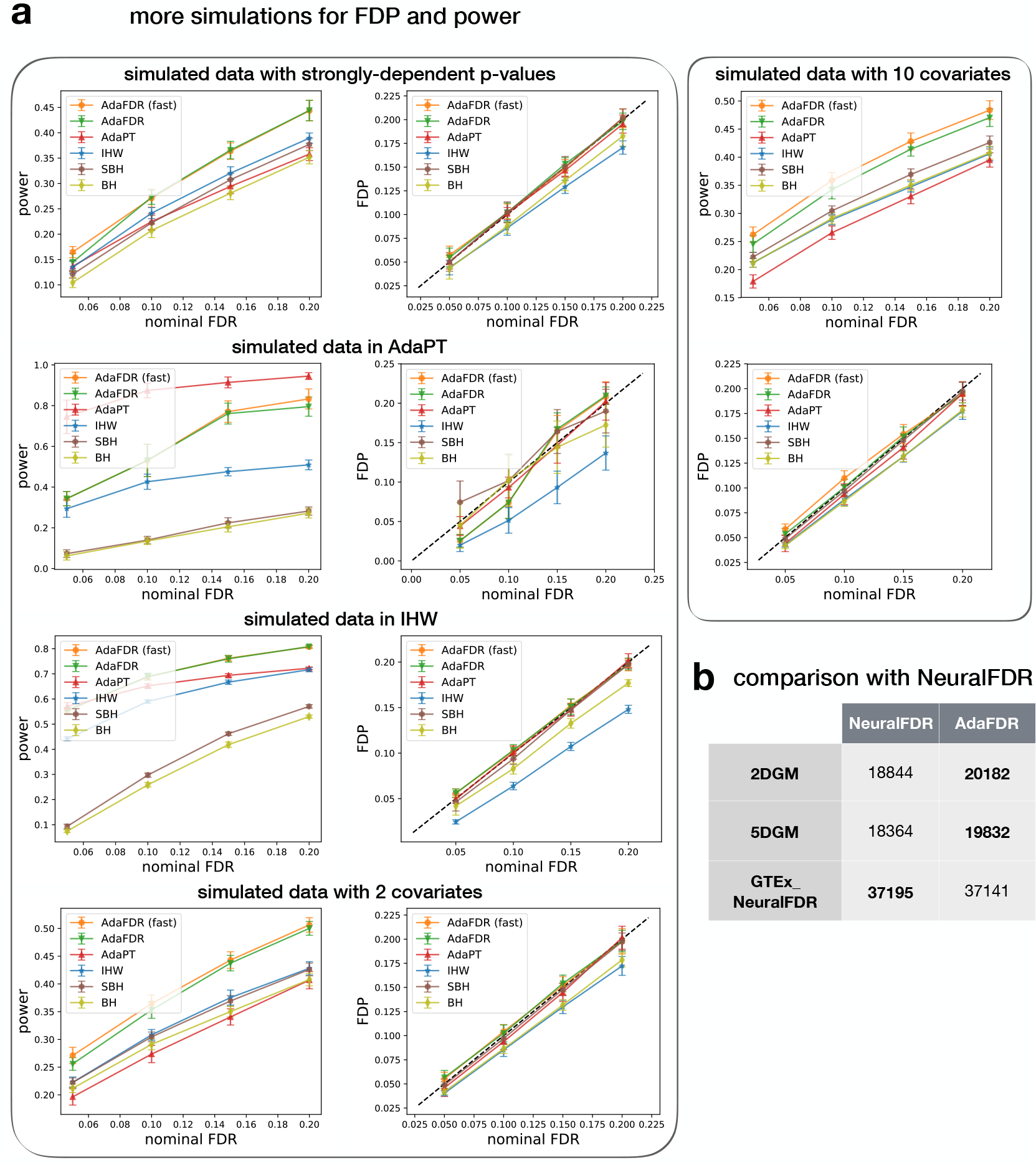
(a) Additional simulations for FDP and power. Descriptions of the data are in Supplementary SubSection 3.6. (b) Comparison between `NeuralFDR` and `AdaFDR`.

**Supplementary Figure 4.**
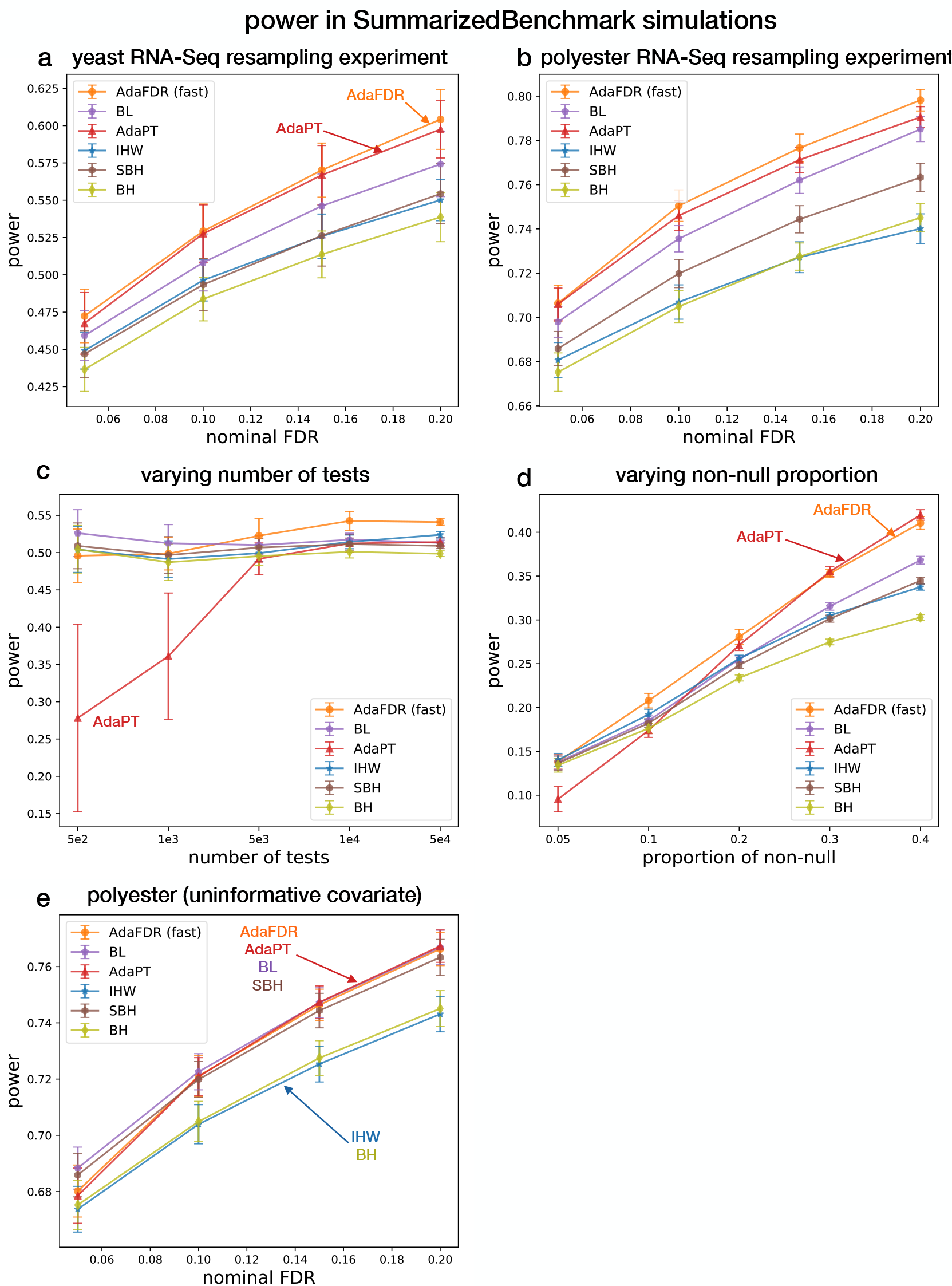
Power in five SummarizedBenchmark simulations^18^ with the corresponding FDP shown in Supplementary Figure 5. Panels a-d correspond to Figure 3 in^18^ while panel e corresponds to the first row of Table S2 in^18^. Performance of an extra method BL^47^ is provided. Performance of `AdaFDR` is very similar to `AdaFDR-fast` and is hence omitted to reduce clutter. Ten resamplings were done for RNA-Seq experiments (a,b,e) while twenty were done for others; 95% confidence intervals are provided. Panels a, b are two RNA-Seq spike-in resampling experiments with an informative covariate, panel c contains a simulated data with the number of tests varying from 500 to 50k, while panel d contains a simulated data with the non-null proportion of tests varying from 0.95 to 0.6. In all four experiments, `AdaFDR` and AdaPT have the highest power (with `AdaFDR` being slightly better). We note that AdaPT does not have such high power in the same experiments in^18^. This is probably because we used adapt_gam while adapt_glm is used in^18^; the former has a better performance but takes a longer time to run. Panel e uses the same set of p-values as panel b but with an uninformative covariate. We can see the performance of IHW reduces to BH while others reduce to SBH, a phenomenon also mentioned in^18^. `AdaFDR` maintains high power here indicating that it is not overfit to the uninformative covariate.

**Supplementary Figure 5.**
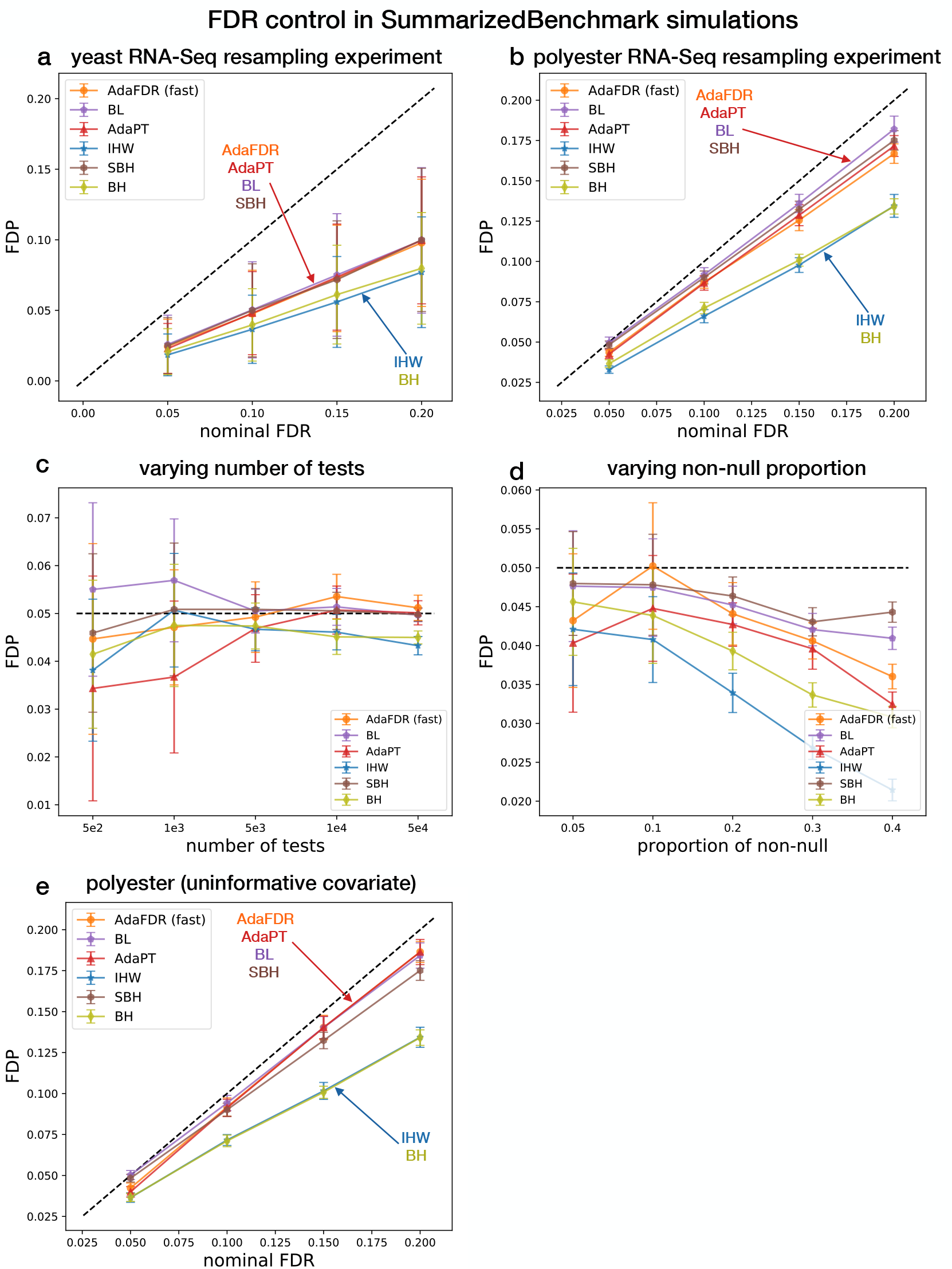
FDR control in five SummarizedBenchmark simulations^18^ with the corresponding power shown in Supplementary Figure 4. The detailed description of the data can also be found in Supplementary Figure 4. Performance of an extra method BL^47^ is provided. Performance of `AdaFDR` is very similar to `AdaFDR-fast` and is hence omitted to reduce clutter. All methods control the FDR accurately, except in panel c, where BL slightly exceeds the FDR control when the number of tests is small.

### 2 Additional Algorithm Information

#### 2.1 Feature preprocessing

We perform feature preprocessing to integrate both numerical covariates and categorical covariates. First for each categorical covariate, the categories are reordered based on the ratio of the alternative probability and the null probability, estimated on the training set using the same method as above. Then quantile normalization is performed for each covariate separately. Note that after this transformation, all covariates will have values between 0 and 1. Also, overfitting is not a concern since the entire proprecessing is done without seeing p-values from the testing set.

#### 2.2 Remark on Theorem 1

Theorem 1 is similar to, but stronger than that for `NeuralFDR`. First, `NeuralFDR` requires the scale factor to be selected from a finite set of *L* numbers and has an extra multiplicative factor 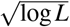 in the error term ε. In contrast, `AdaFDR` selects the scale factor over all positive numbers and the 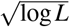 term is gone. This is done by using a stochastic process argument instead of the union bound. Second, `NeuralFDR` uses an empirical Bayes model where the tuples (*P_i_*, **x**_*i*_, *h_i_*) are generated i.i.d. following some hierarchical model. `AdaFDR`, however, requires a less restrictive assumption made only on the conditional distribution of null p-values, whereas the covariates and alternative p-values can have arbitrary dependence.

#### 2.3 Initialization via EM algorithm

Here we present the EM algorithm that is used to fit the mixture model (2) on a set of *N* points 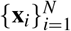. Recall that due to quantile normalization, the value of **x**_*i*_ is within [0,1]^*d*^. Therefore, each component in the mixture model is truncated to be within [0,1]^*d*^, i.e. truncated GLM or truncated Gaussian. Since we need to use the samples each associated with a sample weight, let us consider the general case where each sample **x**_*i*_ receives a positive weight *v_i_* ∈ ℝ_+_.

For the sake of convenience, let us reparameterize the parameters to have the standard probability distribution

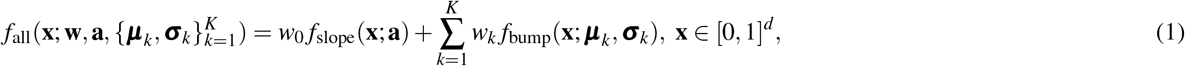

where **w** ∈ [0,1]^*K* + 1^ with 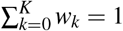 and

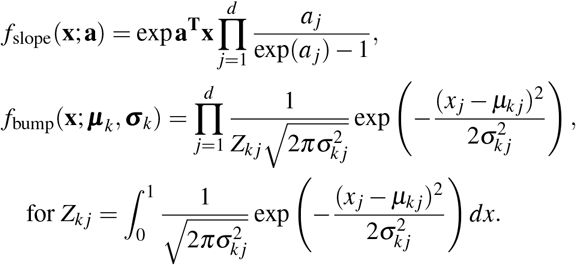

It is not hard to see that (1) is equivalent to the mixture threshold (2) up to a scale factor that can be specified by *b* in (2); knowing one, the parameters for the other can be computed without difficulty.

The EM algorithm can be described as follows. For the initialization, the responsibility **r**_*i*_ ∈ [0,1]^*K*+1^, *i* ∈ [*N*] for each point **x**_*i*_ is initialized as

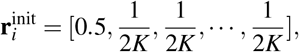

where the first component corresponds to the slope component and the rest correspond to the *K* bump components. Then, the algorithm iterates between the E-step and the M-step as follows until convergence:

1. **Expection (E-step)**: For each point **x**_*i*_, update the responsibility

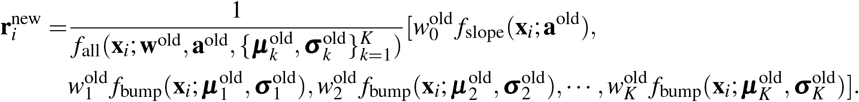
2. **Maximization (M-step)**: Update the component weights **w**^new^ by

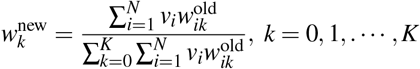

Update the parameters for the slope component and each of the *K* bump component:

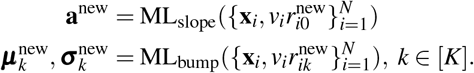

The ML estimates of slope and bump, i.e. 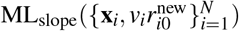 and 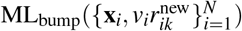, are described as follows.

##### ML estimate of the slope

The log likelihood function of a single observation **x**_*i*_ can be written as

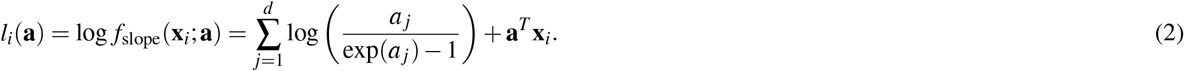

Further the weighted average log likelihood function,

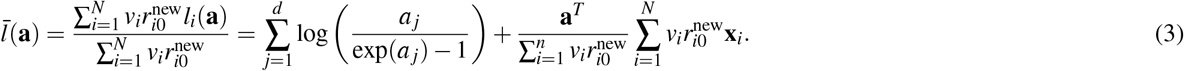

We add a regularization term 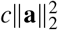 to encourage small values of 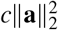, i.e.

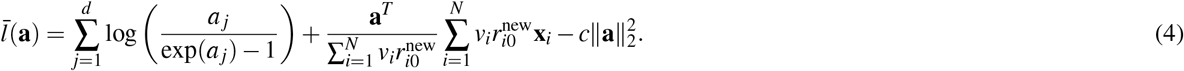

We found that setting *c* = 0.005 gives a stable result. We solve the ML estimation problem by setting the derivative to be zero. Namely, for the *j*-th element *a_j_*,

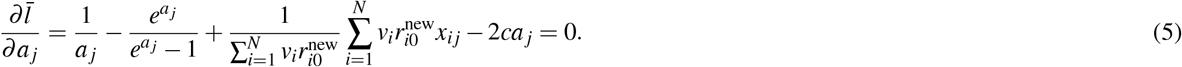

Rearranging terms on both sides we have that the ML estimate *â*_*j*_ satisfies

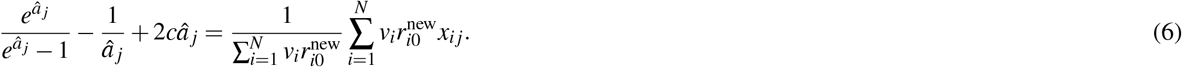

Since the left-hand-side term is monotonic in *â*_*j*_, the ML solution *â*_*j*_ can be computed via binary search.

##### ML estimate of the *k*-th bump

Since the density function can be factorized as a product of different dimensions, the ML estimation can be done for each dimension separately. Now consider observation **x**_*i*_. The log likelihood function corresponding to dimension *j* can be written as

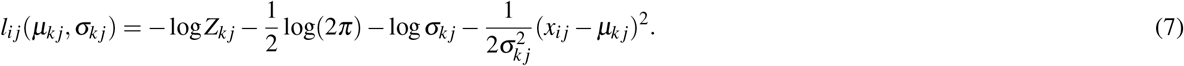

Then the weighted average log likelihood function for dimension *j* can be written as

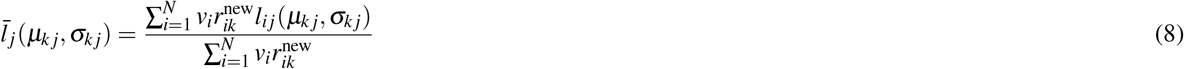

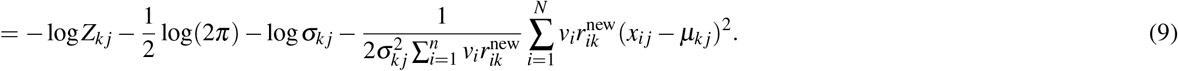

Since 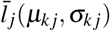 is convex, we compute the ML estimation 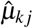 and 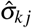 via gradient descent, where the derivatives are given as follows.

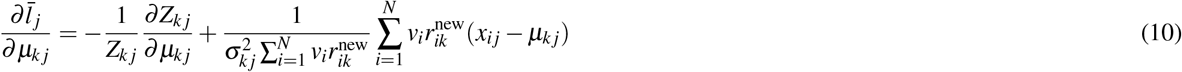

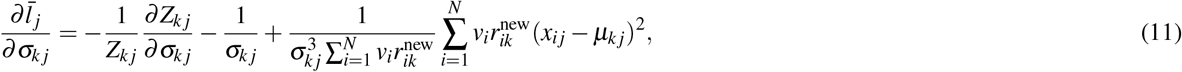

where the derivatives with respect to *Z_kj_* are

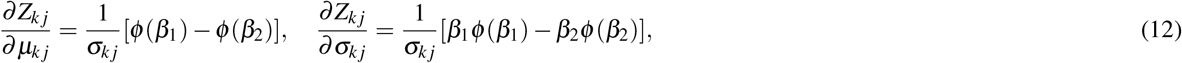

and 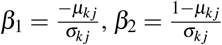 and 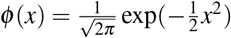.

#### 2.4 Implementation of other methods

1. AdaPT: adapt_gam is used with a 5-degree spline for each dimension. This choice is based on a discussion with the authors of AdaPT.
2. IHW: The covaraites are first clustered into 20 clusters using Kmeans clustering. Then IHW is run with the default setting and the cluster labels as the univariate covariate. This automatically incorporates the multi-dimensional case. For the univariate case, this does not change the result much as compared to directly running IHW. For example, for the airway data, directly running IHW gives 4873 discoveries while Kmeans+IHW gives 4862 discoveries.
3. BL: First the null distribution *π*_0_(x) is estimated using lm_pi0 with 5 degrees of freedom. Then BH is used with p-values weighted by 1/*π*_0_(**x**_*i*_). This is the same as the usage in^18^.

### 3 Data

#### 3.1 eQTL study

##### GTEx

For eQTL study, we used Genotype-Tissue Expression (GTEx) datasets^7^. This datasets aims at characterizing variation in gene expression levels across individuals and diverse tissues of the human body. We used the V7 release of GTEx analysis data (dbGaP Accession phs000424.v7.p2). The datasets contains 11688 samples, and in total there are 53 tissues from 714 donors (44 of them with sample size >70 are used in the GTEx paper). We filtered the tissues based on the following criteria. First, the tissue needs to have eQTL analysis, where the number of samples with genotype is greater than 70. Second, we set the number of samples threshold to be 100 in order to make the p-values more reliable. Third, we would like the tissue to have a corresponding roadmap^8^ cell type, so that we can leverage the cell-specific chromatin state data from roadmap. After filtering, we were left with 17 cell types. The meta-information of the filtered GTEx datasets is listed in Table 1.

**Table 1.**
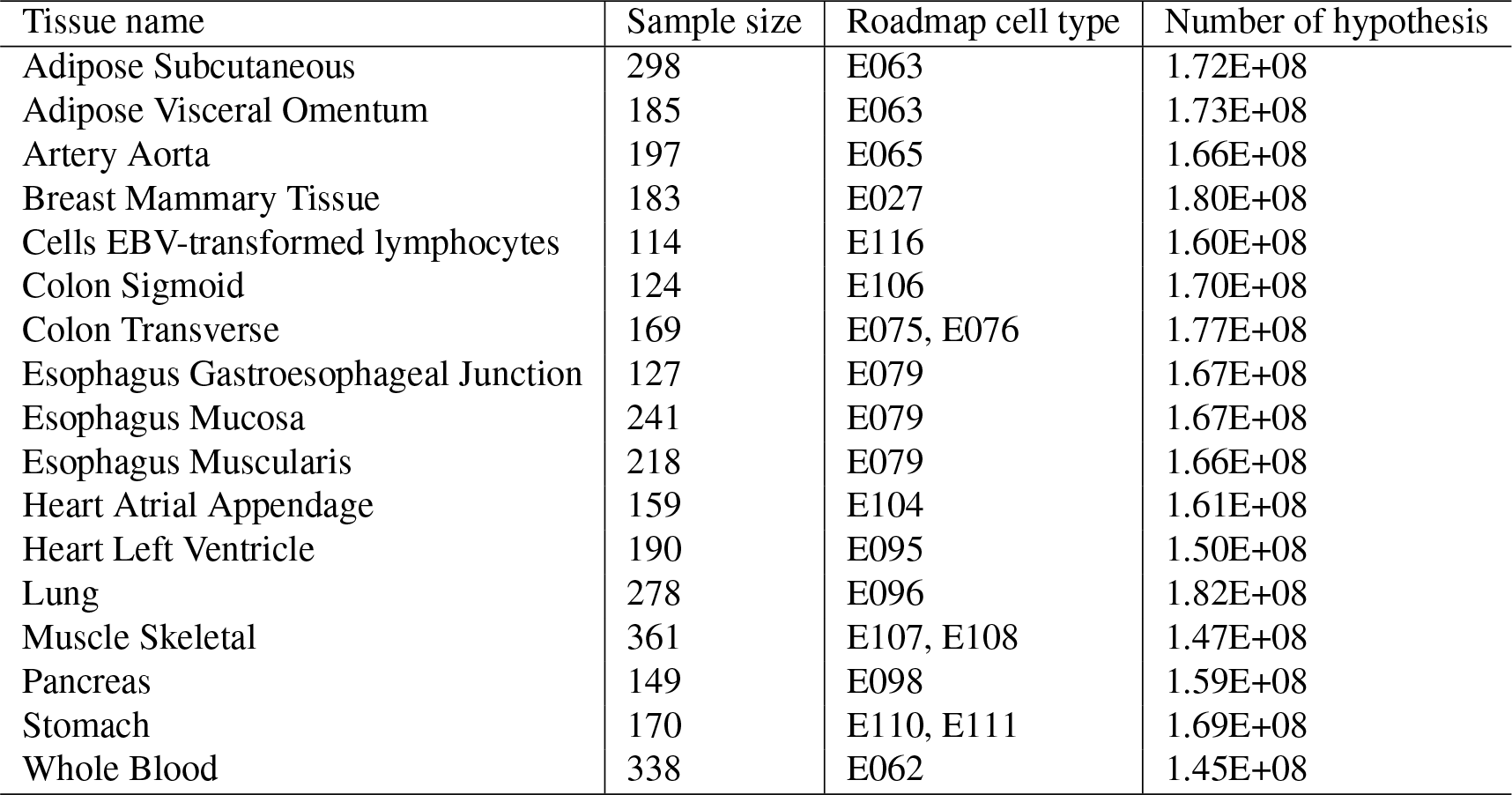
Information for selected GTEx tissue types

| Tissue name | Sample size | Roadmap cell type | Number of hypothesis |
| --- | --- | --- | --- |
| Adipose Subcutaneous | 298 | E063 | 1.72E+08 |
| Adipose Visceral Omentum | 185 | E063 | 1.73E+08 |
| Artery Aorta | 197 | E065 | 1.66E+08 |
| Breast Mammary Tissue | 183 | E027 | 1.80E+08 |
| Cells EBV-transformed lymphocytes | 114 | E116 | 1.60E+08 |
| Colon Sigmoid | 124 | E106 | 1.70E+08 |
| Colon Transverse | 169 | E075, E076 | 1.77E+08 |
| Esophagus Gastroesophageal Junction | 127 | E079 | 1.67E+08 |
| Esophagus Mucosa | 241 | E079 | 1.67E+08 |
| Esophagus Muscularis | 218 | E079 | 1.66E+08 |
| Heart Atrial Appendage | 159 | E104 | 1.61E+08 |
| Heart Left Ventricle | 190 | E095 | 1.50E+08 |
| Lung | 278 | E096 | 1.82E+08 |
| Muscle Skeletal | 361 | E107, E108 | 1.47E+08 |
| Pancreas | 149 | E098 | 1.59E+08 |
| Stomach | 170 | E110, E111 | 1.69E+08 |
| Whole Blood | 338 | E062 | 1.45E+08 |

In this filtered datasets, each hypothesis is a gene-variant pair. Nominal P values for each gene-variant pair were estimated using a two-tailed t-test. Each gene-variant is associated with 4 or 5 covariates listed below:

- **gene expression** We obtained the median gene expression from the gene in gene-variant pair and used as a feature.
- **alternative allele frequency** We mapped each SNP to the dbSNP database^51^. We took the alternative allele frequency as a feature. If there were multiple alternative alleles, we took the smallest one. For the SNPs we cannot find a mapping, this feature is imputed with mean alternative allele frequency.
- **TSS distance** The distance from the SNP to the transcription starting site is used as a feature. It is defined as *pos_SNP_* — *pos_TSS_*.
- **cell-specific chromatin state** We took the position of the SNP and mapped it to roadmap database. Each SNP falls into the 15-state chromatin model. This state is used as a categorical feature.
- **p-value from another tissue (optional)** Optionally, we used the P value from another tissue as a covariate. If we cannot find the same gene-variant pair in another tissue, we impute with the mean P value. This covariate is only used for “AdaFDR (aug)” and “AdaFDR (ctrl)” experiments.

##### MuTHER

In the Multiple Tissus Human Expression Resource project^46^, samples from 850 individuals were collected and 3 tissues, namely adipose, LCL, and skin, were studied. We used only the data for the adipose tissue, where a nominal p-value is provided for each SNP-gene pair.

#### 3.2 RNA-Seq data

We used three RNA-Seq datasets to validate our algorithm. The first one `bottomly`^15^ is an RNA-Seq datasets used to detect differential gene expression between mouse strains. We used the same data preprocessing pipeline as in IHW^12^. p-values were calculated using `DESeq2`, and the mean of normalized counts for each gene were chosen to be the covariate. The second datasets `airway`^14^ is an RNA-Seq datasets used to identify the differentially expressed genes in airway smooth muscle cell lines in response to dexamethasone. The datasets is processed with the same pipeline as `bottomly`. The thrid datasets `Pasilla`^52^ is an RNA-Seq datasets for detecting genes that are differentially expressed between the normal and Pasilla-knockdown conditions. This datasets is available in `Pasilla` package and it is analyzed in the vignette of `genefilter` package using independent filtering method. The p-values were generated using `DESeq` package and the logarithm of normalized count were used as the covariate. All the preprocessing steps can be reproduced using vignettes of R package `IHWPaper`^53^.

#### 3.3 Microbiome data

The two microbiome experiments are from the benchmark paper^18^.

#### 3.4 Proteomics data

The proteomics data is from the IHW paper^12^.

#### 3.5 fMRI data

The two fMRI data are from the fMRI paper^20^.

#### 3.6 Simulated data

##### Data 1. Simulated data with one covariate

The covariate **x**_*i*_ ~ Unif[0,1] and the probability of being an alternative hypothesis given the covariate ℙ(*h*_*i*_ = 1|**x**_*i*_) is defined using the mixture model (1) as

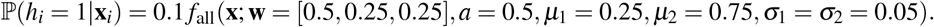

The null p-values are generated i.i.d. from Unif[0,1] while the alternative p-values are generated i.i.d. from *P*_*i*_ ~ Beta(α = 0.3, *β* = 4). The number of hypotheses is 20000 and 10 datasets are generated with different random seeds.

##### Data 2. Simulated data with two covariates

The covariate **x**_*i*_ ~ Unif[0,1] and the probability of being an alternative hypothesis given the covariate ℙ(*h*_*i*_ = 1|**x**_*i*_) is defined using the mixture model (1) as

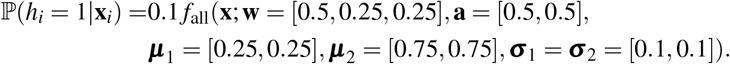

The null p-values are generated i.i.d. from Unif[0,1] while the alternative p-values are generated i.i.d. from *P*_*i*_ ~ Beta(α = 0.3, *β* = 4). The number of hypotheses is 20000 and 10 datasets are generated with different random seeds.

##### Data 3. Simulated data with ten covariates

First a simulated data with two covariates is generated (data 2). Then, another 8 noisy dimensions are added to the covariates with each entry drawn i.i.d. from Unif[0,1]. The number of hypotheses is 20000 and 10 datasets are generated with different random seeds.

##### Data 4. Simulated data with weakly-dependent p-values

The covariate **x**_*i*_ ~ Unif[0,1] and the probability of being an alternative hypothesis given the covariate ℙ(*h*_*i*_ = 1|**x**_*i*_) is generated same as the simulated data with one covariate (data 1). The p-values are converted to z-scores via *p* = 1 - Φ(*z*), where Φ(.) is the cdf of the standard normal distribution. Every 10 consecutive null z-scores are generated from 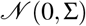, while every 10 consecutive alternative z-scores are generated from 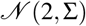, with the symmetric covariance matrix whose upper triangular part is specified as

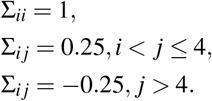

We note instead of 0.25, the value 0.4 is used in the original paper (Section 3.2,^4^). However such choice makes the covariance matrix not positive semi-definite. We decrease the value until the matrix becomes positive semi-definite. The number of hypotheses is 20000 and 10 datasets are generated with different random seeds.

##### Data 5. Simulated data with strongly-dependent p-values

The setting is the same as the weakly dependent data (data 4) except the generation of z-scores. Here, every 5 consecutive null z-scores are generated from 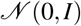, while every 5 consecutive alternative z-scores are generated from 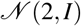. This perfect correlation means to model the linkage disequilibrium (LD) that frequently occurs in SNPs. Due to the reduction of the inherent multiplicity, the number of hypotheses is increased to 50000. 10 datasets are generated with different random seeds.

##### Data 6. Simulated data used in AdaPT

The same data for Figure 6a in^24^ is used where the number of hypotheses is 2500. 10 datasets are generated with different random seeds.

##### Data 7. Simulated data used in IHW

The data is generated according to Supplementary Section 4.2.2 in^12^ where the number of hypotheses is 20000. While the original paper varies the effect size from 1 to 2.5 (the shift of z-scores for alternative p-values), here we only use a fixed effect size 2. 10 datasets are generated with different random seeds.

### 4 Proofs and Auxiliary Lemmas

#### 4.1 Proof of Theorem 1

##### Proof

To avoid ambiguity, we make a few clarifications before the proof. First, the entire analysis is done while conditioning on the hypothesis splitting, all covariates {**x**_*i*_}_*i*∈[*N*]_, the type of hypotheses {*h*_*i*_}_*i*∈[*N*]_, and the alternative p-values 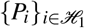, hence allowing arbitrary dependencies of them. Here we note that the reason for splitting the hypotheses at random is to attain good power. The randomness of the analysis comes from the null p-values, which are assumed to be i.i.d. uniformly distributed for convenience. A discussion on extending to the case where the null p-values, conditional on the covariates, are independently distributed and stochastically greater than the uniform distribution is provided at the end.

We also clarify a few notations. We use 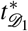 to denote the threshold which is learned on fold 1 and will be applied on fold 2. 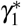 denotes the scale factor of fold 1. For the testing-related quantities, we use subscript “1” to denote those evaluated on fold 1, including the number of discoveries 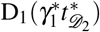, the number of false discoveries 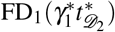, the mirror-estimated number of false discoveries 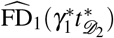 and the mirror-estimated false discovery proportion 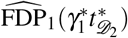. Note that here 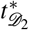 is the threshold that is learned on fold 2 and then applied on fold 1. The term inside the bracket, 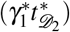, may be omitted when there is no concern of being ambiguous. Quantities for fold 2 are defined in a similar fashion. Now we preceed to the proof.

###### Step 1: show that in order to prove the result, it suffices to show that

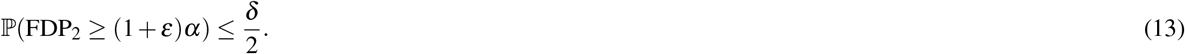

Indeed, if (13) it true, then by symmetry 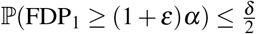. Further by the union bound, with probability (w.p.) at least 1-δ,

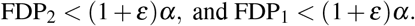

This further implies that w.p. at least 1-*δ*, the FDP on the whole datasets

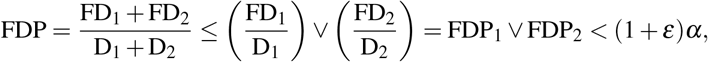

which gives the desired result. Hence in the rest of the proof, we denote effort to proving (13). Also, since we are only to deal with fold 2, we drop the subscript 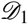 for threshold learned on fold 1 to have 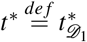.

###### Step 2: convert the probability ℙ(FDP_2_ ≥ (1 + ε)α) to some analyzable stochastic process

Let 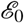 denote the set of random variables that we wish to condition on, including hypothesis splitting, all covariates {**x**_*i*_}_*i*∈[*N*]_, the type of hypotheses {*h*_*i*_}_*i*∈[*N*]_, and the alternative p-values 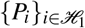. Let us consider the conditional version of (13):

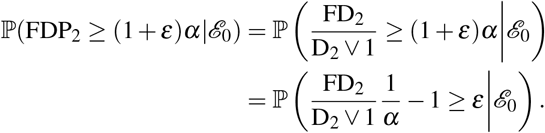

Let 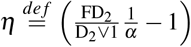. Recall that FD_2_ and D_2_ correspond to the best rescaled threshold on fold 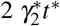 and the best scale factor 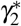 is selected from the set 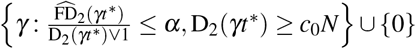. Then *η* can be upper bounded by

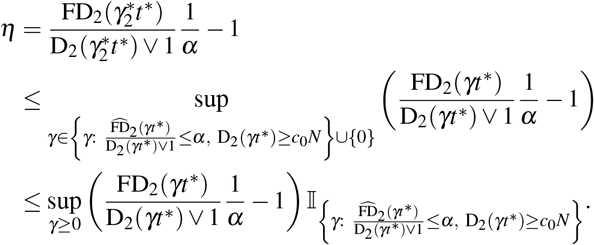

Furthermore, since the indicator function is one only when 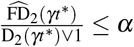, which can also be written as 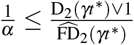 with the convention that 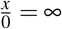 for any *x* > 0, we further have

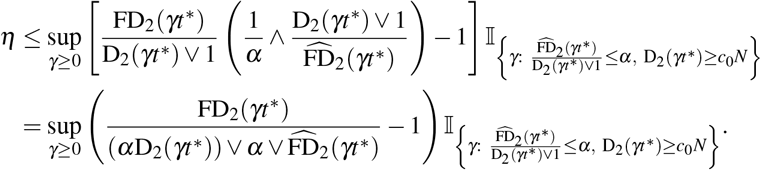

Again since indicator function is one only when D_2_(γ*t*\*) > *c*_0_*N*,

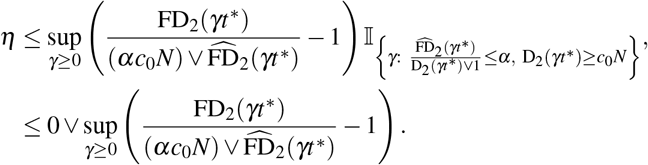

Furthermore with the notation 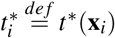 where we recall that we have defined 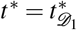 before,

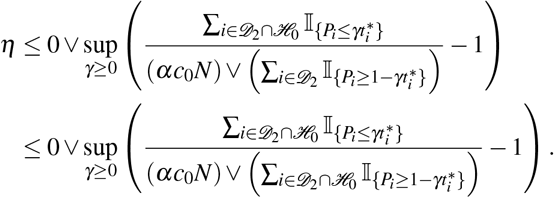

Finally, we can complete the conversion by noting that

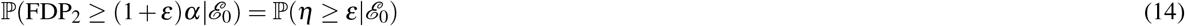

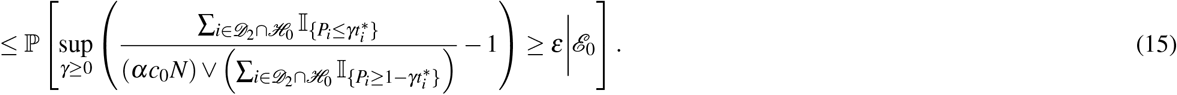

Here, the first term in (15), i.e. 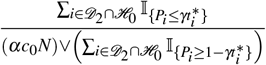, can be understood as a stochastic process that as *γ* grows from 0 to infinity, new elements are added to the numerator and the denominator with equal probability. Hence this term should always be close to 1. We next proceed to prove the result following this intuition.

###### Step 3: Upper bound the probability of (15)

We note that the p-values involved in (15) are all null p-values from fold 2. Hence, they are i.i.d. uniformly distributed conditional on 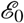. Let 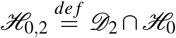. For any *i* ∈ ℋ_0,2_, *γ* > 0, define the random variables

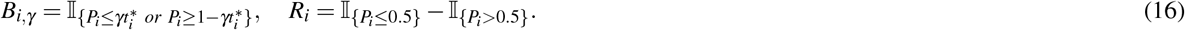

Since ∀*i* ∈ ℋ_0,2_, 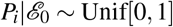, we have 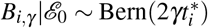 and 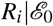 are i.i.d. Rademacher random variables. In addition, it is easy to verify that *B*_*i,γ*_ is independent of *R_i_* and

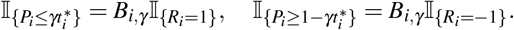

Hence (15) can be written in terms of *B*_*i,γ*_’s and *R_i_*’s as

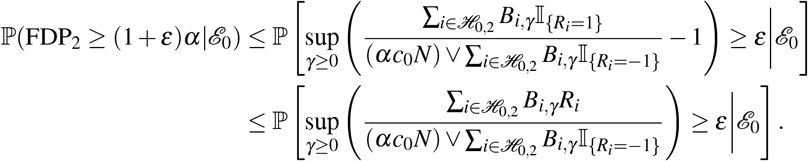

Furthermore, let 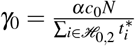. Divide the set of γ in the sup from [0, ∞) into [0, γ_0_] and (γ_0_, ∞), and apply union bound to have

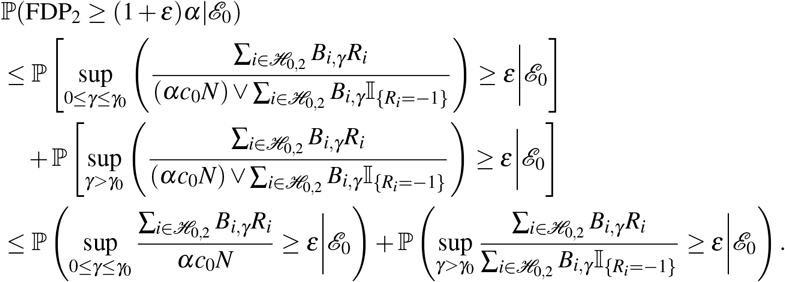

Define the random set 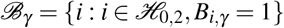. We note that the sequence of sets 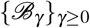 is monotonic in the sense that as γ grows, more elements are incorporated into 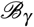. With this definition, the above inequality can be further written as

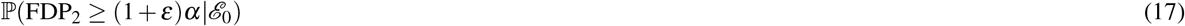

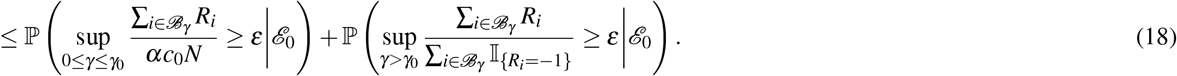

Next we upper bound the two terms in (18) respectively. Here let us use 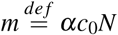 for simplicity.

###### The first term in (18)

For some *m*_0_ > 2*m* to be specified later, by the law of total probability,

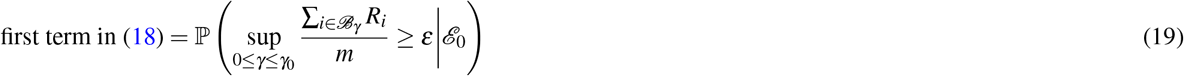

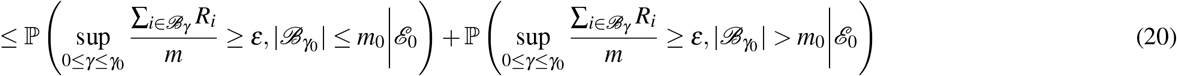

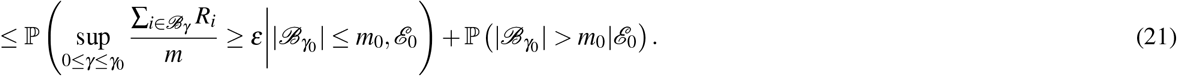

The two terms in (21) are upper bounded separately. Consider the first term. Recall that 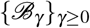 is a random sequence of monotonic sets; let 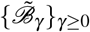 denote any of its realization. Then since taking expectation over all possible 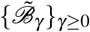 s.t. 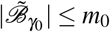 is no greater than taking the sup of them,

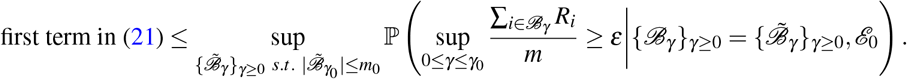

Consider the term inside the probability, i.e. 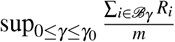, where due to conditioning 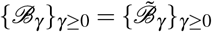. Recall that the sequence 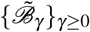 is monotonic that as γ grows more elements are incorporated into the set but no element is removed from the set. Also up to the point γ = γ_0_ there are altogether 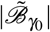 elements. Then the sup is equivalent to being evaluated over a sequence of 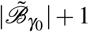 monotonic sets, i.e. 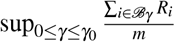 is equal to 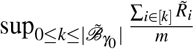 in distribution, where 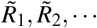 is a sequence of i.i.d. Rademacher random variables independent of everything else. Therefore,

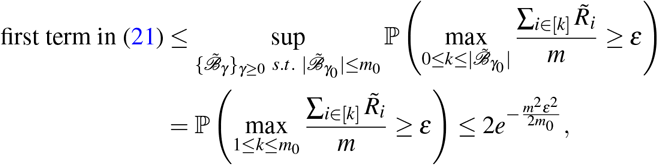

where the last inequality is due to Lemma 1.

Now consider the second term in (21). Recall that 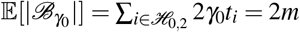 by the definition of γ_0_. By Lemma 2,

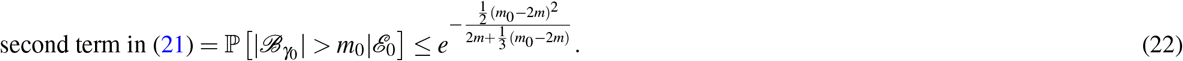

By setting *m*_0_ = 3*m*, we have

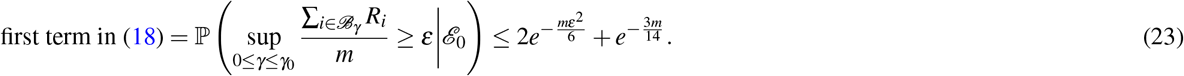

###### The second term in (18)

For some *m*_1_ ≤ 2*m* to be specified later, by the law of total probability,

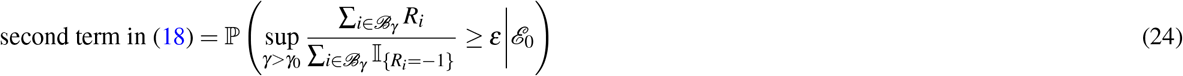

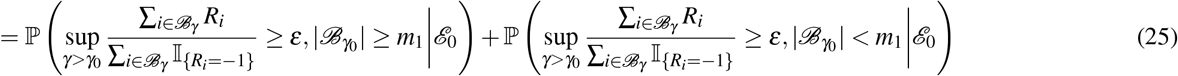

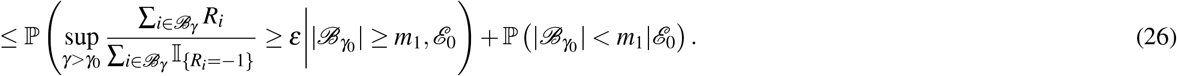

Using the same argument for analyzing the first term in (21),

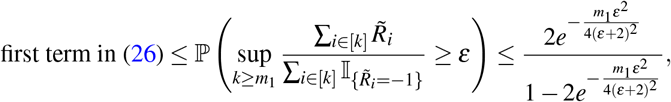

where we recall that 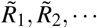 is a sequence of i.i.d. Rademacher random variables indepedent of everything else, and the second inequality is due to Lemma 1.

Similar to (22), by Lemma 2,

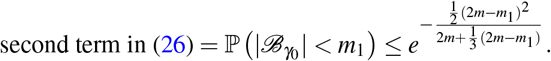

By setting *m*_1_ = *m*, we have

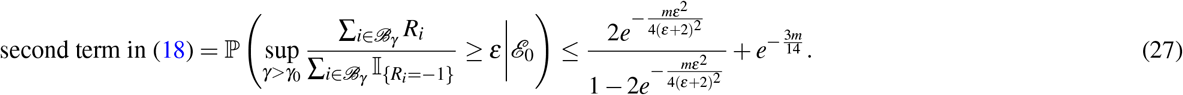

Combining (23) and (27) we have that for (18),

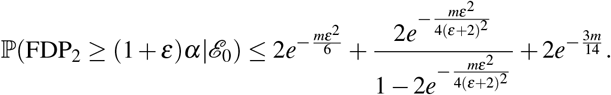

Furthermore,

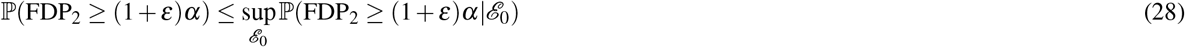

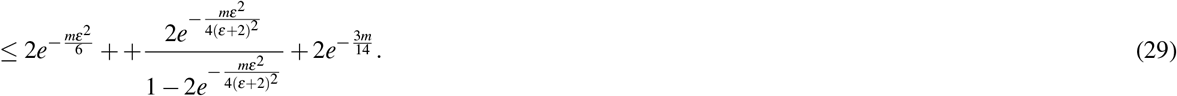

By equaling the term in the right-hand-side of (29) with 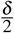 we have 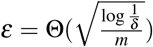. Recall that *m* = *αc*_0_*N* where *c*_0_ is a constant, we have

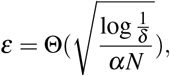

which concludes the proof.

In order for the proof to hold, it is required that the mirror estimate 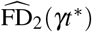 is stochastically no less than the true number of false discoveries FD_2_(*γt*\*) for any *γ* ≥ 0. This is still true when the i.i.d. assumption for the null p-values is extended to the assumption that the null p-values, conditional on the covariates, are independently distributed and stochastically greater than the uniform distribution. Hence the result is directly extendable.

#### 4.2 Lemma 1 with proof

##### Lemma 1.

*(Some properties of random walk) Let R*_1_,*R*_2_, … *be i.i.d. Rademacher random variables and let 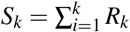. Then for any integer n* > 1 *and for any real number t* > 0,

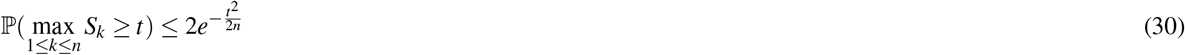

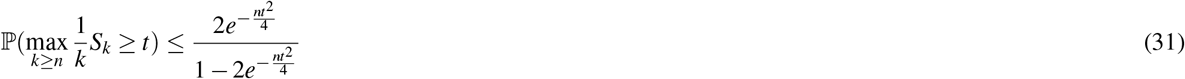

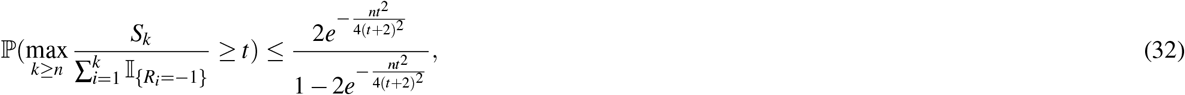

*where for the second and the third inequalities, we require t to be large enough for the probability to be positive*.

*Proof*. (30) is proved via a standard reflection argument for random walk. First consider when *t* is an integer,

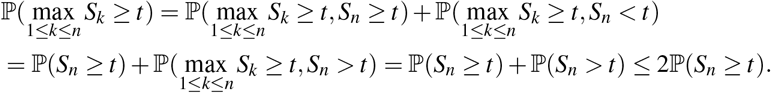

If *t* is not an integer,

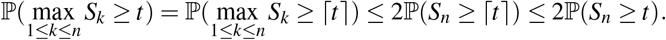

Finally, using Hoeffding’s inequality, one has

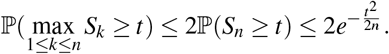

(31) is proved via a technique called “peeling”. Specifically,

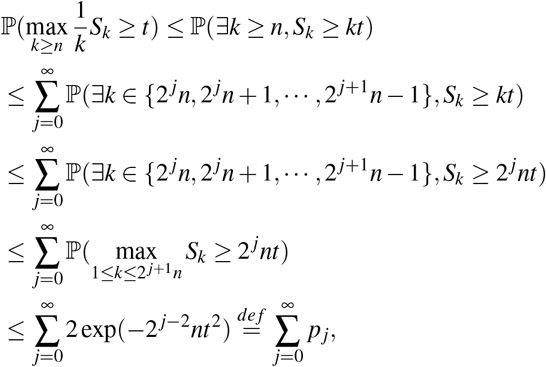

where the last inequality is due to (30) that we have just proved. Note that for 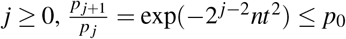. Hence,

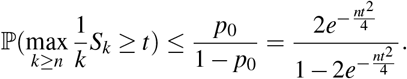

Finally, (32) is a direct consequence of (31):

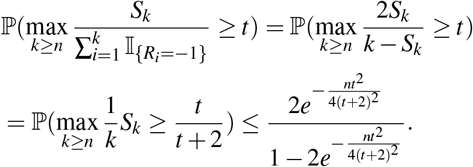

#### 4.3 Lemma 2 with proof

##### Lemma 2

*(Some properties ofnon-homogeneous Bernoulli sum) Let B_i_ ~ Bern(p_i_) be some independent Bernoulli random variables. Then*

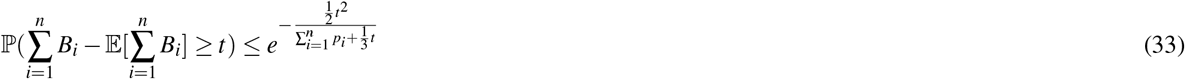

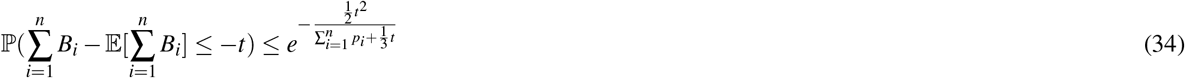

*Proof*. Define 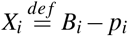. Then *X_i_*’s have zero means and are independent of each other. Also, note that |*X_i_*| ≤ 1 almost surely and 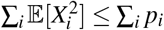. Hence (33) and (34) can be obtained by applying Bernstein inequality on {*X_i_*} and {−*X_i_*} respectively.

